# Cytosolic factors controlling PASTA kinase-dependent ReoM phosphorylation

**DOI:** 10.1101/2024.04.02.587704

**Authors:** Patricia Rothe, Sabrina Wamp, Lisa Rosemeyer, Jeanine Rismondo, Joerg Doellinger, Angelika Gründling, Sven Halbedel

## Abstract

Bacteria adapt the biosynthesis of their envelopes to specific growth conditions and prevailing stress factors. Peptidoglycan (PG) is the major component of the cell wall in Gram-positive bacteria, where PASTA kinases play a central role in PG biosynthesis regulation. Despite their importance for growth, cell division and antibiotic resistance, the mechanisms of PASTA kinase activation are not fully understood. ReoM, a recently discovered cytosolic phosphoprotein, is one of the main substrates of the PASTA kinase PrkA in the Gram-positive human pathogen *Listeria monocytogenes*. Depending on its phosphorylation, ReoM controls proteolytic stability of MurA, the first enzyme in the PG biosynthesis pathway. The late cell division protein GpsB has been implicated in PASTA kinase signalling. Consistently, we show that *L. monocytogenes prkA* and *gpsB* mutants phenocopied each other. Analysis of *in vivo* ReoM phosphorylation confirmed GpsB as an activator of PrkA leading to the description of structural features in GpsB that are important for kinase activation. We further show that ReoM phosphorylation is growth-phase dependent and that this kinetic is reliant on the protein phosphatase PrpC. ReoM phosphorylation was inhibited in mutants with defects in MurA degradation, leading to the discovery that artificial MurA overexpression prevented ReoM phosphorylation. Overexpressed MurA must adopt its substrate-bound closed conformation and interact with ReoM to exert this effect, but the extracellular PASTA domains of PrkA or MurJ flippases where not required. Our results indicate that intracellular signals control ReoM phosphorylation and extend current models describing the mechanisms of PASTA kinase activation.

## Introduction

The main component of bacterial cell walls is peptidoglycan (PG), a network of glycan strands that are connected with each other by short peptide bridges. This mesh engulfs the cell and serves as a protective layer against external influences but also acts as a mechanical antagonist of cellular turgor. PG constitutes up to ∼30% of the dry weight of a Gram-positive bacterial cell and therefore requires a high amount of precursor molecules and energy equivalents for its biosynthesis. Due to these massive energy costs, PG biosynthesis is closely coordinated with growth and nutrient supply and can also be activated in response to PG damage (1–3). Biosynthesis of PG starts in the cytoplasm with the consumption of UDP-*N*-acetylglucosamine (UDP-GlcNAc) by the MurA and MurB enzymes that sequentially build up UDP-*N*-acetylmuramic acid (UDP-MurNAc). Further reactions assemble a pentapeptide side chain at the MurNAc unit, transfer the resulting molecule onto the lipid carrier undecaprenyl phosphate and further add a second GlcNAc moiety (4, 5). The resulting lipid linked pentapeptide-disaccharide (called lipid II) is then flipped across the membrane by so-called flippases (6, 7). On the extracellular side, the disaccharide unit is transferred onto growing PG chains by glycosyltransferases and later crosslinked by transpeptidases. These latter two enzymatic activities are either provided by bifunctional class A penicillin binding proteins (PBPs) or by SEDS-type glycosyltransferases that cooperate with class B PBPs that are mere transpeptidases (8–11). The PG network is very dynamic and also remodelled by degradative enzymes to adjust size and shape to growth, division and development (12). PG biosynthesis is fuelled either with UDP-GlcNAc generated by the GlmSMU enzymes from fructose-6-phosphate, an intermediate of glycolysis (4), or by salvage of PG precursors from environmental sources (13).

PG biosynthesis is controlled at multiple steps in the pathway and by different regulatory mechanisms. This includes transcriptional activation of biosynthetic and PG remodelling genes by alternative sigma factors and two component systems (3, 14, 15), regulation of *glmS* translation through a metabolite sensitive riboswitch (16, 17), control of MurA protein stability (18–20), negative feedback inhibition of enzyme activities by downstream intermediates (21–23), activation of PBPs and their recruitment into protein complexes by interaction with scaffolding proteins (24–26) and enzyme phosphorylation by PASTA domain containing serine/threonine protein kinases (27–29).

We and others have described a central PASTA kinase-dependent mechanism of PG biosynthesis regulation. This mechanism controls the proteolytic degradation of MurA by the ClpCP protease (19, 20, 30, 31). MurA, which catalyses the first committed step in the PG biosynthesis pathway, is a known ClpCP substrate in *Bacillus subtilis*, *Listeria monocytogenes*, *Enterococcus faecalis* and *Staphylococcus aureus* (18–20, 32). MurA degradation is highly regulated and requires the assistance of three additional proteins: ReoM (IreB in *E. faecalis*), ReoY and MurZ (also known as MurAB or MurA2) (19, 20, 30, 33). One of these, ReoM, is phosphorylated at a conserved threonine residue (Thr-7 in *L. monocytogenes* ReoM) by its cognate PASTA kinase, called PrkA in *L. monocytogenes* (19, 30, 34). In the unphosphorylated form, ReoM directly interacts with MurA and hands it over to ClpCP for degradation (19, 20, 35). However, upon phosphorylation of ReoM at Thr-7 by PrkA, P∼ReoM cannot bind MurA any longer, which then accumulates (19, 20, 36). The roles of ReoY and MurZ in this process are not well understood, but bacterial two hybrid data suggest that ReoY binds ClpCP as well as ReoM and therefore likely acts as a bridge between both (19). MurZ is a paralogue of MurA, but cannot take over its function in *B. subtilis* and *L. monocytogenes* (18, 33), whereas the two MurA homologues of *E. faecalis*, *S. aureus* and *Streptococcus pneumoniae* can at least partially replace each other (37–39).

MurA enzymes consist of two globular domains and undergo conformational changes upon substrate binding. While the interdomain cleft is open in unliganded MurA, substrate binding induces a movement of the two domains towards each other so that liganded MurA adopts a more compact conformation (40). Substrate binding was required for IreB (the *E. faecalis* homologue of ReoM) to interact with MurAA from *E. faecalis* (20), suggesting that ReoM only binds MurA in its closed conformation.

ReoM phosphorylation is reversed by the protein phosphatase PrpC (19), however, neither the mechanisms controlling PrkA or PrpC activity are fully understood. PASTA kinases of *B. subtilis*, *E. faecalis* and *S. pneumoniae* are activated by GpsB, a late cell division protein with transient septal localisation (41–43). PASTA kinases have further been shown to bind lipid II and/or muropeptides through their extracellular PASTA domains (44–46). When PknB, the PASTA kinase of *Mycobacterium tuberculosis*, is modified in a way such that it can no longer bind lipid II, the protein becomes hyperactive (46), suggesting an inhibitory role of lipid II.

Here, we investigated the regulation of ReoM phosphorylation in *L. monocytogenes*. For this, we established a method to directly measure ReoM phosphorylation *in vivo* using a ReoM specific antibody and separation of differently phosphorylated ReoM species by native polyacrylamide gel electrophoresis.

## Results

### *L. monocytogenes* mutants in *gpsB* and *prkA* phenocopy each other

We previously demonstrated that a *L. monocytogenes* Δ*gpsB* mutant cannot grow at increased temperature (24). However, this growth defect was suppressed by mutations in *clpC*, *reoY*, *reoM*, *murA*, *murZ* and *prpC* (19, 33, 35). Interestingly, a similar spectrum of suppressor mutations was reported by Kelliher *et al.* to correct the hypersensitivity of a conditional *prkA* mutant against ceftriaxone (30) (Fig. 1A). This overlap in the spectrum of suppressor mutations suggested that GpsB and PrkA could act in the same cellular pathway(s) and – as a consequence – that inactivation of their genes results in similar phenotypes. The *prkA* gene is essential in *L. monocytogenes* strain EGD-e (19), but deletion of the C-terminal PASTA domains is tolerated and generates a partial *prkA* phenotype (47). To test the assumed phenotypic similarity, we compared the ceftriaxone sensitivity of *gpsB* and *prkA* mutants. This confirmed increased susceptibility of a *L. monocytogenes* strain depleted for PrkA and of a *prkA* mutant lacking the PASTA domains (*prkA*ΔC) against ceftriaxone. Remarkably, the *gpsB* mutant was also hypersensitive against ceftriaxone (Fig. 1B), as was reported recently in another *L. monocytogenes* strain background (48). Furthermore, we also found that the *prkA*ΔC mutant showed a pronounced growth defect at 42°C (Fig. 1C-D), as is known for the *gpsB* mutant (24). Another characteristic phenotype of the Δ*gpsB* mutant emerges when *divIVA* is additionally deleted in the same strain: Δ*gpsB* Δ*divIVA* double mutant cells are considerably longer compared to either single mutant (24). Interestingly, a similar elongation of cells was observed in a *prkA*ΔC Δ*divIVA* double mutant (Fig. 1E). Moreover, the *prkA*ΔC mutant was less prone to lysis when treated with lysozyme as compared to wild type, and a similar phenotype had been reported for a Δ*gpsB* mutant (49) and was confirmed here (Fig. 1F). Taken together, this shows that mutants in *prkA* and *gpsB* generally exhibit similar phenotypes, reinforcing the idea that PrkA and GpsB cooperate.

**Figure 1:**
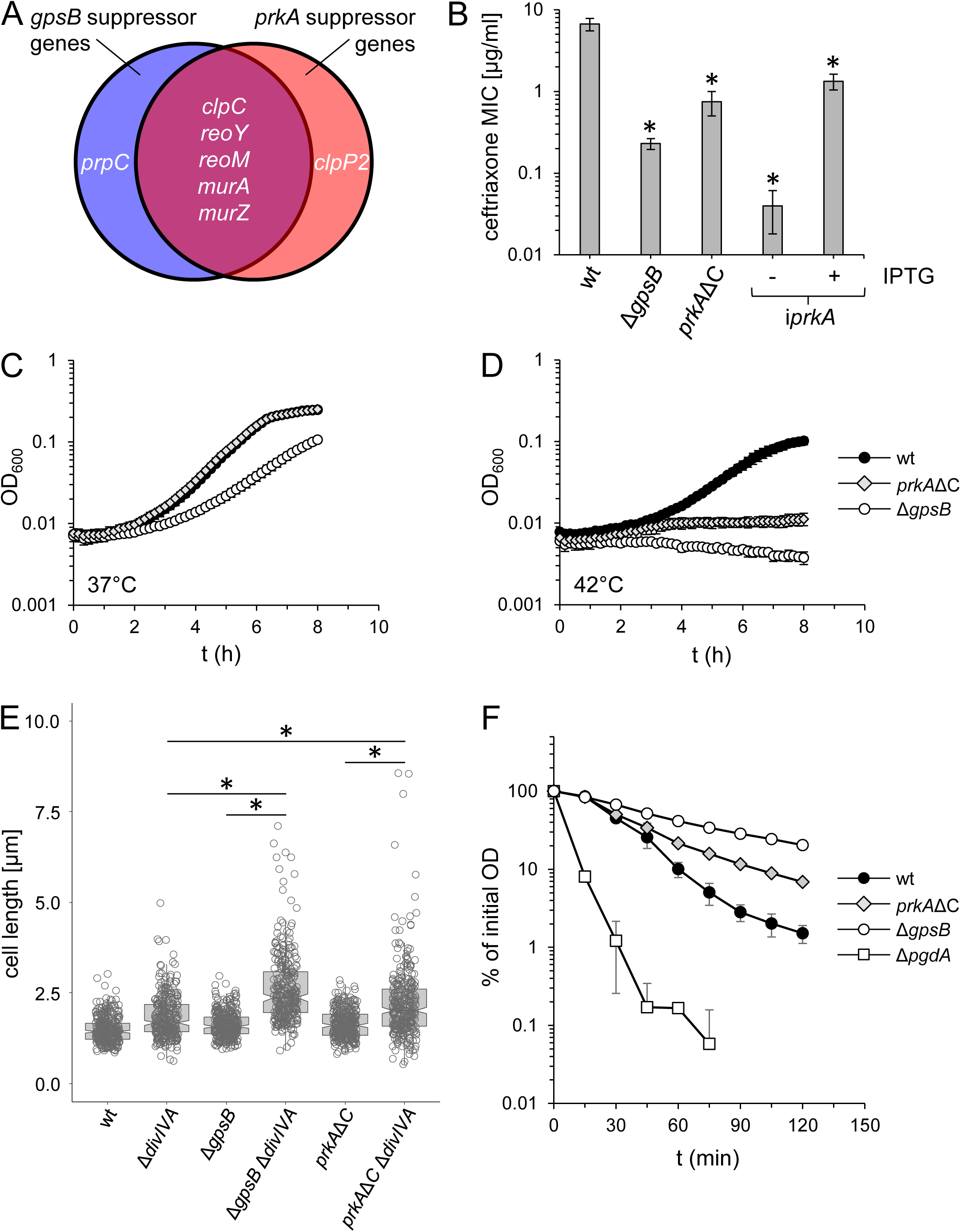
Phenotypic similarity of *L. monocytogenes gpsB* and *prkA* mutants. (A) Mutants in *gpsB* and *prkA* are suppressed by mutations in the same set of genes. Venn diagram showing genes that acquired mutations in suppressors of a *L. monocytogenes* Δ*gpsB* mutant (EGD-e background) (19, 33, 35) and in suppressors of a conditional *prkA* mutant (10403S background) (30). (B) Mutants in *gpsB* and *prkA* are hypersensitive to ceftriaxone. Minimal inhibitory concentrations of ceftriaxone for *L. monocytogenes* strains EGD-e (wt), LMJR19 (Δ*gpsB*), LMS278 (*prkA*ΔC) and LMSW84 (i*prkA*) grown ± IPTG. The experiment was repeated three times and average values and standard deviations are shown. Asterisks indicate statistically significant differences compared to wild type (*P*<0.01, *t*-test with Bonferroni-Holm correction). (C-D) *gpsB* and *prkA* mutants cannot grow at 42°C. Growth of *L. monocytogenes* strains EGD-e (wt), LMJR19 (Δ*gpsB*) and LMS278 (*prkA*ΔC) in BHI broth at 37°C (C) and 42°C (D). (E) Introduction of the Δ*divIVA* deletion into *gpsB* and *prkA* mutant strains results in strong cell elongation. Box plot showing cell lengths of *L. monocytogenes* strains EGD-e (wt), LMS2 (Δ*divIVA*), LMJR19 (Δ*gpsB*), LMJR28 (Δ*gpsB* Δ*divIVA*), LMS278 (*prkA*ΔC) and LMPR27 (*prkA*ΔC Δ*divIVA*) during mid-logarithmic growth in BHI broth at 37°C. 300 cells per strain were measured. Asterisks indicate significance levels (*P*<0.001, *t*-test with Bonferroni-Holm correction). (F) Mutants in *gpsB* and *prkA* are less susceptible to lysozyme. *L. monocytogenes* strains EGD-e (wt), LMJR19 (Δ*gpsB*), LMS278 (*prkA*ΔC) and LMS163 (Δ*pgdA*) were challenged with lysozyme and decline of optical density over time was recorded. Average values and standard deviations from three technical replicates are shown.

### *In vivo* PrkA activity depends on GpsB

Given the phenotypic similarities of *gpsB* and *prkA* mutants described above, it was reasonable to assume that *L. monocytogenes* PrkA requires GpsB for activity, as reported for other Gr+ bacteria (41–43). Previously it has been reported that P∼ReoM can be separated from unphosphorylated ReoM using native PAGE in *in vitro* experiments (19). To use this technique for separation of the two ReoM species in cellular extracts, an antiserum was generated against *L. monocytogenes* ReoM. During initial experiments, this antiserum did only poorly detect ReoM in wild type extracts. We thus introduced a *reoM-his* allele under control of the strong P*_help_* promoter to facilitate ReoM detection (50). This allowed detection of a single ReoM-His species in the wild type (Fig. 2A). For comparison with cells lacking PrkA, we inserted the *reoM-his* allele into the Δ*prkA murA N197D* background, where lethality of the Δ*prkA* deletion is overcome by the *murA N197D* mutation (35). ReoM-His migrated slower in this strain (Fig. 2A), which is in good agreement with the retarded migration of unphosphorylated ReoM observed *in vitro* (19). In contrast to the wild type, two ReoM species were detected in the *murA N197D* background, one at the same height as in wild type and an additional one, the origin of which is currently unclear, but probably corresponds to a ReoM/ReoM-P heterodimer (Fig. 2A). Next, we tried to introduce the *reoM-his* allele into the Δ*gpsB* mutant, but consistent with the idea that GpsB is required for PrkA activity and with the toxicity of unphosphorylated ReoM, this was never possible. Therefore, *reoM-his* was brought into the Δ*gpsB murAN197D* background, yielding a viable strain. ReoM-His migrated at the position of the unphosphorylated ReoM species observed in the Δ*prkA murAN197D* strain (Fig. 2A), supporting the idea that GpsB is required for PrkA activity in *L. monocytogenes*.

**Figure 2:**
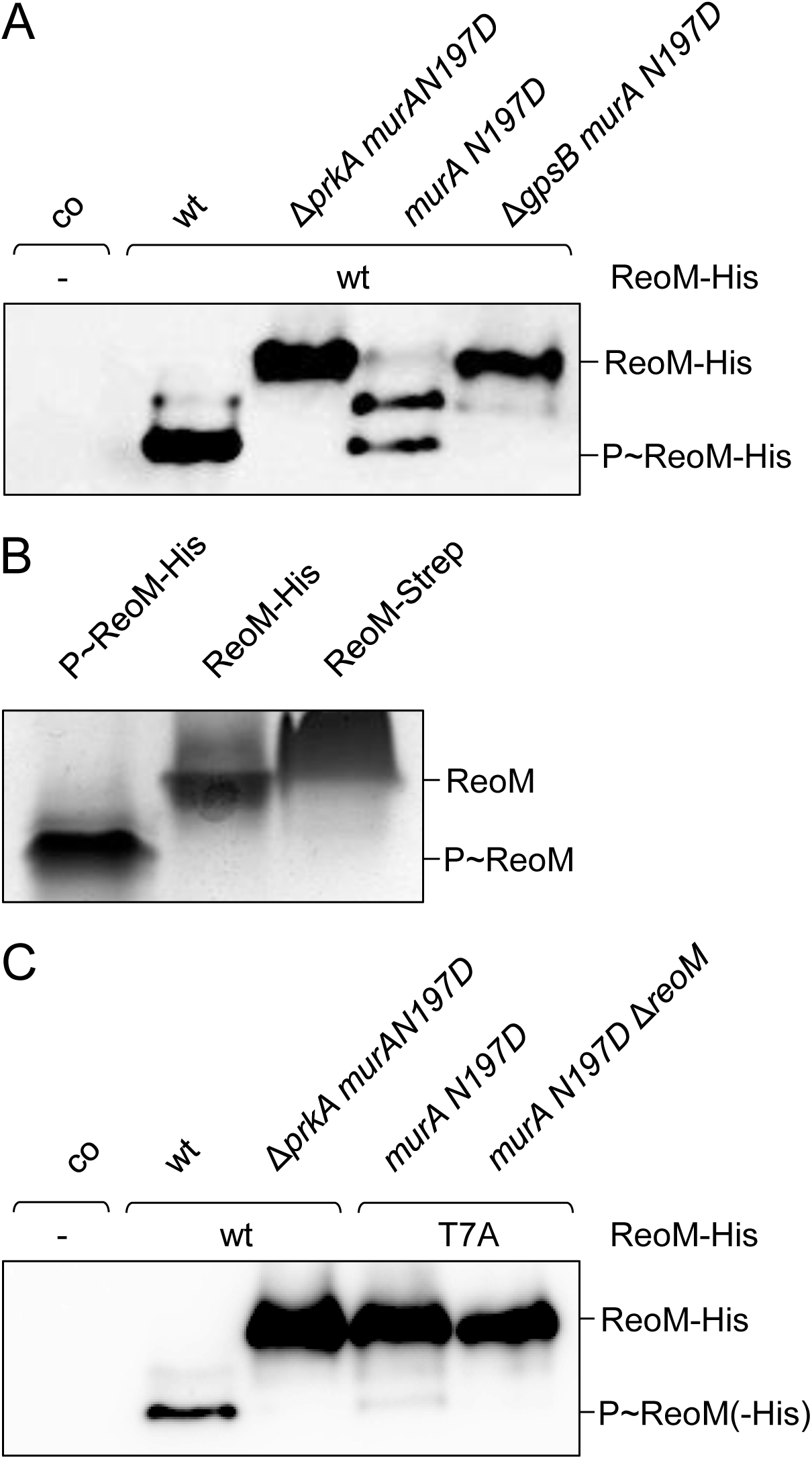
GpsB is required for *L. monocytogenes* PrkA activity. (A) Western blot after native PAGE for detection and separation of the different forms of ReoM-His in *L. monocytogenes* strains LMJD22 (labelled “wt”), LMPR5 (“Δ*prkA murA N197D”*), LMPR9 (“*murA N197D”*) and LMPR10 (“Δ*gpsB murA N197D”*). *L. monocytogenes* EGD-e (“co”) was included as negative control. The presence of ectopically expressed ReoM-His is specified. (B) Native PAGE loaded with ReoM-His∼P purified from strain LMJD22 (*prkA^+^*) and with unphosphorylated ReoM-His, which was purified from the Δ*prkA murA N197D* strain LMPR5. ReoM-Strep directly purified from *E. coli* was included as control. (C) *In vivo* ReoM phosphorylation requires Thr-7. Western blot after native PAGE for ReoM detection in *L. monocytogenes* strains EGD-e (labelled “co”), LMJD22 (“wt”), LMPR5 (“Δ*prkA murA N197D”*), LMPR42 (*“murA N197D*”) and LMPR48 (“*murA N197D* Δ*reoM”*). The presence of ectopically expressed ReoM-His and T7A variants is specified.

We then purified the two ReoM-His species from the two *L. monocytogenes* strains LMJD22 (*prkA*^+^) and LMPR5 (Δ*prkA murA N197D*) (Fig. 2B). ReoM-His purified from the *prkA*^+^ strain run faster through the native gel than ReoM-His purified from the Δ*prkA murA N197D* strain, the position of which was similar to unphosphorylated ReoM-Strep purified from *E. coli* (Fig. 2B). The two ReoM-His species were subjected to mass spectrometry, which showed phosphorylation at Thr-7 and Tyr-10 in ReoM purified from wild type background (LMJD22), while ReoM purified from Δ*prkA* cells (LMPR5) was only phosphorylated at Tyr-10.

As final proof that we look at ReoM phosphorylated at Thr-7 here, we analysed the *in vivo* ReoM phosphorylation patterns in a strain expressing a phospho-ablative *reoM T7A-his* variant as second *reoM* copy (LMPR43). As expected, ReoM T7A-His migrated at the position of the unphosphorylated ReoM in this strain (Fig. 2C), however a faint signal at the position of the phosphorylated protein remained, which corresponded to the native endogenous ReoM, as this signal was no longer detected upon *reoM* deletion (strain LMPR48, Fig. 2C).

### ReoM phosphorylation is generally insensitive to defects in cell division

GpsB is a late cell division protein (51) and PASTA kinases are known to localize to cell division sites in *B. subtilis* and *S. pneumoniae* (52, 53). We therefore wondered, whether cell division proteins other than GpsB also influence PrkA activity and analysed phosphorylation of endogenous ReoM, which became detectable by loading larger amounts of protein, in strains lacking the non-essential cell division genes *divIVA*, *pbpA1*, *sepF*, *zapA*, *minCD* and *minJ*. However, ReoM was found to be phosphorylated in all these mutants except the Δ*gpsB* strain (Fig. S1A), demonstrating that the effect of GpsB on PrkA was specific.

Next, we asked whether distortion of cell division by depletion of an essential cell division protein would influence ReoM phosphorylation. PBP B2 is required for septal PG biosynthesis and its depletion blocks cell division leading to formation of long non-septated filaments (54). We depleted PBP B2 in strain LMJR18 and analysed phosphorylation of endogenous ReoM by Western blotting. However, no effects on ReoM phosphorylation were found (Fig. S1B). Likewise, depletion of PBP B1 (strain LMJR27), necessary for PG biosynthesis along the lateral cell cylinder and maintenance of rod shape (54), did not interfere with phosphorylation of endogenous ReoM (Fig. S1B). Apparently, ReoM phosphorylation is not disturbed by defects in cell division in general and also does not depend on the two specific modes of PG biosynthesis at the septum and the lateral wall.

### ReoM phosphorylation requires hexameric GpsB

GpsB is a hexameric membrane protein known to interact with PBP A1 through a defined cleft in its surface (24, 25, 55). Moreover, GpsB itself is phosphorylated by PrkA at various threonine residues including Thr-88 (30, 42, 55). To address the question which functionalities in GpsB support its role in PrkA activation, we determined the ReoM phosphorylation pattern in strains expressing *gpsB* alleles with specific functional mutations. As can be seen in Fig. S2, wild type-like ReoM phosphorylation patterns were observed in *gpsB* L24A and R24A mutants. These mutations interfere with binding of GpsB to the cytoplasmic membrane (24). Likewise, normal ReoM phosphorylation was detected in strains carrying *gpsB* alleles with mutations preventing PBP A1 binding (Y27A-I40A) (25) (Fig. S2). Remarkably, normal ReoM phosphorylation was also observed in a strain, in which *gpsB* carried the phospho-ablative T88A mutation. When the same residue was mutated in a phospho-mimetic manner (T88D), a slight reduction in ReoM phosphorylation became apparent (Fig. S2). In contrast, strong effects on ReoM phosphorylation were observed in *gpsB* mutants, in which mutations prevented the interaction of the two trimeric C-terminal domains required for hexamer formation (F91A, L94A, F105A) (55) or when even the trimers could not be formed due to mutations in residues essential for formation of the trimer stabilizing salt bridges (R96A, E101A) (24) (Fig. S2). This shows that formation of the hexameric GpsB complex is of great importance for PrkA activation, while GpsB does not need to be membrane-bound or bound to PBP A1 to exert this effect. Likewise, phosphorylation of GpsB at Thr-88 seems to play only a minor role for activation of PrkA compared to the structural effects.

### ReoM phosphorylation changes with growth progression

Next, ReoM phosphorylation was studied during different stages of growth. For this, a culture of strain EGD-e was grown in BHI broth at 37°C and samples were taken at different optical densities. ReoM was found to be phosphorylated during exponential growth phase but a shift towards unphosphorylated ReoM was detected during stationary phase (Fig. 3A). We also observed that MurA levels declined during growth and were reduced down to 13±6% at late stationary phase (OD_600_=3.0) compared to the original level detected at an OD_600_ of 0.5 (Fig. 3A). This demonstrates that ReoM phosphorylation and MurA levels decline at stationary phase.

**Figure 3:**
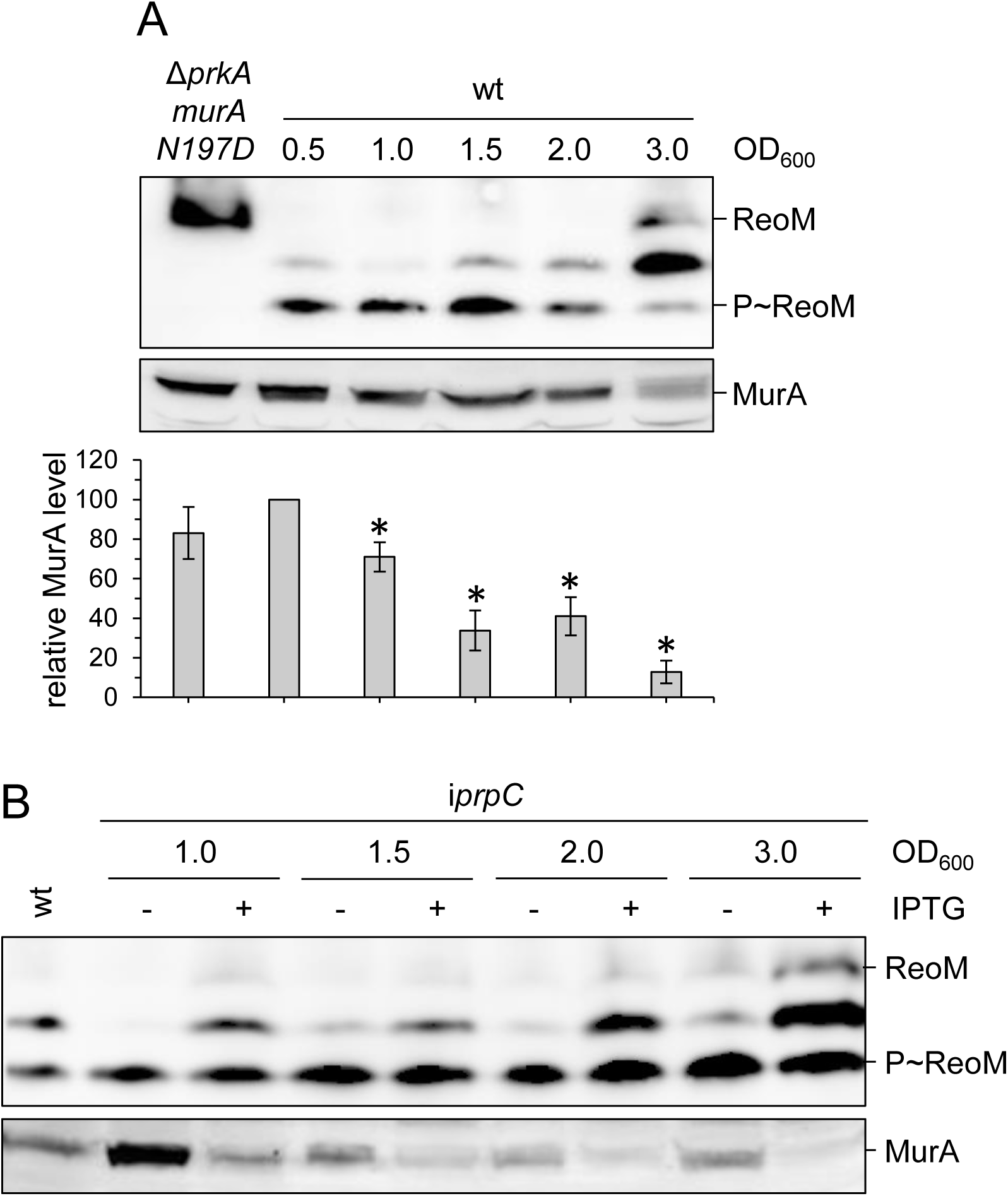
Effect of growth phase on ReoM phosphorylation. (A) Growth phase dependent ReoM phosphorylation and MurA levels. Western blot after native PAGE showing ReoM phosphorylation in *L. monocytogenes* strain EGD-e (wt) that was grown in BHI broth at 37°C to the indicated optical densities (upper panel). Strain LMS266 (Δ*prkA murA N197D*) was included as negative control. Western blot after standard SDS PAGE to determine MurA levels in the same samples (middle panel). MurA signals were determined by densitometry and expressed as values relative to wild type at OD_600_=0.5 (bottom panel). Average values and standard deviations are shown (n=3) and asterisks label statistically significant differences relative to wild type at OD_600_=0.5 (*P*<0.05, *t*-test with Bonferroni-Holm correction). (B) Decline of ReoM phosphorylation and MurA levels in stationary phase depends on PrpC. Western blot after native PAGE showing ReoM phosphorylation in *L. monocytogenes* strain EGD-e (wt) and LMSW83 (i*prpC*) in BHI broth ± 1 mM IPTG at 37°C to the indicated optical densities (upper panel). Western blot showing MurA levels in the same samples (lower panel).

### Control of ReoM phosphorylation by PrkA and PrpC

The antagonist of PrkA is the phosphatase PrpC, which dephosphorylates P∼ReoM *in vitro* (19). The *prpC* gene is essential in wild type (19, 47). To test for PrpC phosphatase activity *in vivo* despite this limitation, we analysed ReoM-His phosphorylation a Δ*gpsB* Δ*prpC* mutant background, where deletion of *prpC* is tolerated (35). To ensure comparability of strains in the intended experiment, we made use of the Δ*gpsB* Δ*prpC* mutant strain LMPR34, which also carried the *murA N197D* mutation. ReoM-His was fully phosphorylated in this strain background, even though no P∼ReoM was found in its progenitor strain LMPR10 (Δ*gpsB murA N197D*) that still contained *prpC* (Fig. S3A). Thus, PrpC dephosphorylates ReoM *in vivo*, which in turn implies that PrkA must still have residual kinase activity despite the absence of GpsB.

We next addressed the contribution of PrpC to the observed growth phase dependent ReoM phosphorylation pattern and determined ReoM phosphorylation in the i*prpC* strain LMSW83 allowing IPTG-dependent PrpC production. When this strain was grown without IPTG, the decline in ReoM phosphorylation and reduction of MurA levels during late stages of growth were delayed (Fig. 3B), indicating that PrpC has a critical role in the growth phase-dependent ReoM dephosphorylation, particularly during stationary phase.

Similar to *prpC*, the *prkA* gene is essential in EGD-e, but deletion of the PASTA domains was tolerated (47). We therefore wondered how deletion of the PASTA domains would affect ReoM phosphorylation. The ReoM-His migration pattern shifted from the fully phosphorylated to the mono-phosphorylated dimer and the unphosphorylated forms in a *prkA*ΔC mutant lacking all PASTA domains (Fig. S3B). Thus, the PASTA domains are required for full PrkA activity. Next, we investigated whether the intermediate kinase activity observed in the *prkA*ΔC background was still GpsB-dependent and compared ReoM-His phosphorylation in the *prkA*ΔC and the Δ*gpsB prkA*ΔC mutants (both containing the *murA N197D* mutation to ensure viability). Deletion of *gpsB* was dominant over the *prkA*ΔC mutation, as only unphosphorylated ReoM-His was detected in the Δ*gpsB prkA*ΔC double mutant (Fig. S3C). This shows that GpsB exerts its impact on PrkA in a manner that is independent of the PASTA domains.

### ReoM phosphorylation responds to MurA accumulation

ReoM acts in concert with ReoY and MurZ to control MurA degradation via ClpCP (19, 35). We wondered whether any of these proteins would be involved in control of P∼ReoM formation and analysed phosphorylation of endogenous ReoM in Δ*clpC*, Δ*murZ* and Δ*reoY* mutants. Remarkably, most ReoM was found unphosphorylated in each of these mutants (Fig. 4A). MurA strongly accumulates in mutants lacking *clpC*, *murZ* or *reoY* (19, 33) (Fig. 4A), but due to the pleiotropic effect that can be expected upon disrupting ClpCP-dependent proteolysis, it is not known whether MurA accumulation is the underlying reason for the effect on ReoM phosphorylation. To address this, we tested the effect of artificial *murA* overexpression on ReoM phosphorylation in strain LMJR123, carrying an IPTG-inducible copy of *murA*. Addition of IPTG led to strong accumulation of MurA in this strain and to a concomitant reduction of ReoM phosphorylation (Fig. 4B). In contrast to this, ReoM phosphorylation was not affected by overexpression of MurZ (Fig. S4). Furthermore, we observed that depletion of MurA from Δ*clpC* cells partially restored phosphorylation of ReoM (Fig. 4B). This demonstrates that ReoM phosphorylation is sensitive to the level of MurA.

**Figure 4:**
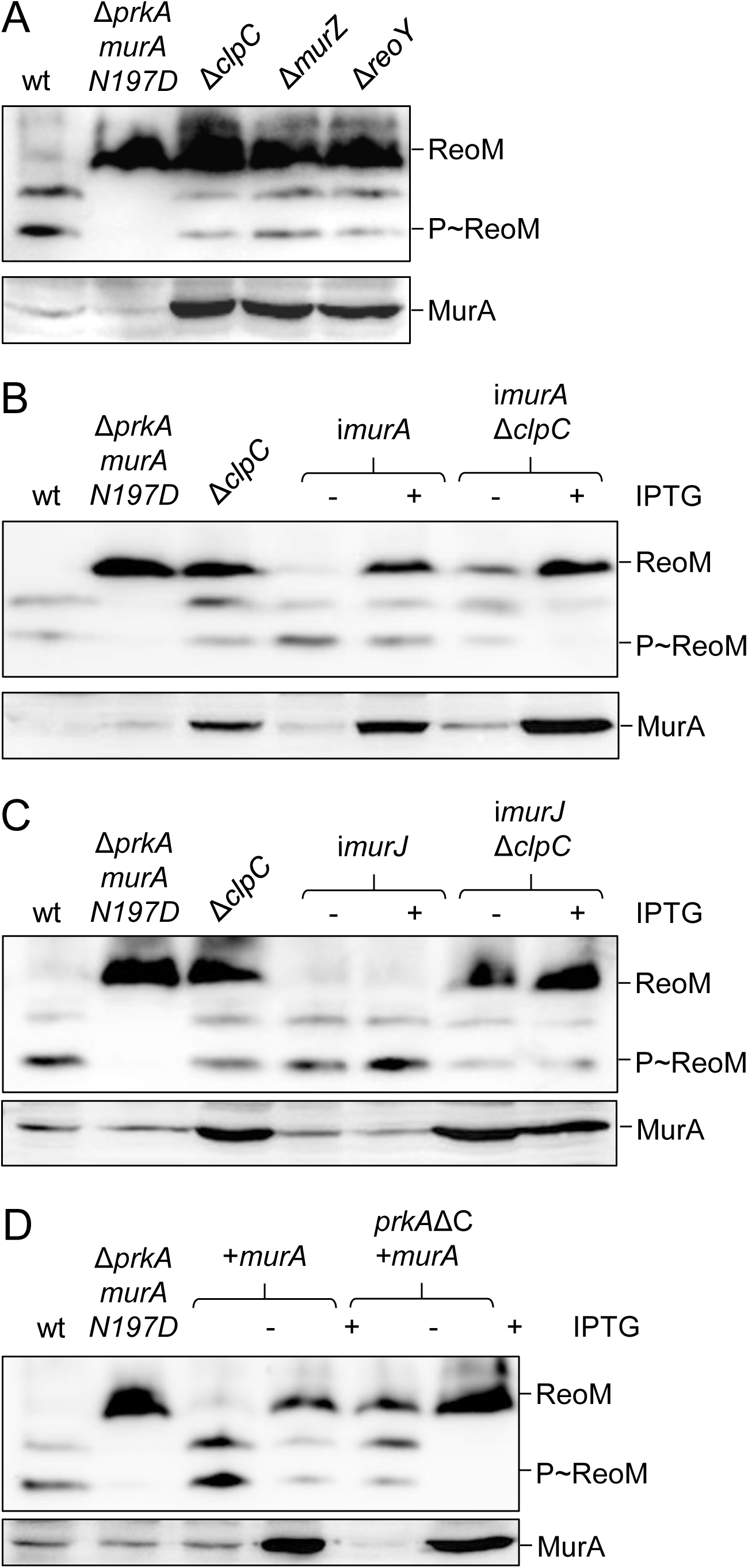
Direct control of ReoM phosphorylation through the level of MurA. (A) ReoM phosphorylation requires ClpC, MurZ and ReoY. Phosphorylation of ReoM was analysed in strains EGD-e (wt), LMS266 (Δ*prkA murA N197D*), LMJR138 (Δ*clpC*), LMJR104 (Δ*murZ*) and LMSW32 (Δ*reoY*) (upper panel). MurA levels in the same set of strains were visualized in a parallel Western blot (lower panel). (B) MurA levels control ReoM phosphorylation in wild type and Δ*clpC* cells. Western blots showing ReoM phosphorylation (upper blot) and MurA levels (bottom blot) in strains EGD-e (wt), LMS266 (Δ*prkA murA N197D*), LMJR138 (Δ*clpC*), LMJR123 (i*murA*) and LMPR52 (i*murA* Δ*clpC*) grown in BHI broth ± 1 mM IPTG. (C) Control of ReoM phosphorylation through MurA is MurJ-independent. Western blots showing ReoM phosphorylation (upper blot) and MurA levels (bottom blot) in strains EGD-e (wt), LMS266 (Δ*prkA murA N197D*), LMJR138 (Δ*clpC*), ANG5140 (i*murJ*) and LMPR54 (i*murJ* Δ*clpC*) grown in BHI broth ± 1 mM IPTG. (D) The MurA effect on ReoM phosphorylation does not involve the PASTA domains of PrkA. Western blots showing ReoM phosphorylation (upper blot) and MurA levels (bottom blot) in strains EGD-e (wt), LMS266 (Δ*prkA murA N197D*), LMJR116 (+*murA*) and LMPR49 (*prkA*ΔC+*murA*) grown in BHI broth ± 1 mM IPTG.

Work in *M. tuberculosis* suggested that PASTA kinases can be inhibited by lipid II through an interaction with the distal PASTA domains (46). Inspired by this result, we assumed that MurA accumulation would lead to higher lipid II levels that in turn could interfere with ReoM phosphorylation by inhibition of PrkA through interaction with its PASTA domains. To test this, we first wanted to determine ReoM phosphorylation in a Δ*clpC* mutant lacking the two MurJ-like lipid II flippases. As simultaneous deletion of both *murJ* genes was not possible, a strain was generated in which an IPTG-dependent copy of *murJ2* (*lmo1625*, *LMRG_01341*) was expressed from an ectopic site and the *murJ2* and *murJ1* (*lmo1624*, *LMRG_01342*) genes were deleted from the chromosome. This strain showed a slight but significant growth retardation (Fig. S5A) and a fourfold reduced resistance to lysozyme in the absence of IPTG (Fig. S5B) indicating at least partial depletion of flippase activity. However, depletion of MurJ activity had no effect on ReoM phosphorylation in these cells (Fig. 4C). Moreover, when MurJ activity was depleted in Δ*clpC* cells, ReoM phosphorylation was not restored (Fig. 4C) as observed with depletion of MurA (Fig. 4B). Apparently, prevention of lipid II transport across the cytoplasmic membrane does not interfere with ReoM phosphorylation. To test this idea further, the effect of MurA overexpression on P∼ReoM formation was determined in *prkA*ΔC cells lacking the PrkA PASTA domains. In these cells, MurA overexpression still exerted a negative effect on ReoM phosphorylation despite the absence of the PASTA domains (Fig. 4D). Thus, the PASTA domains of the *L. monocytogenes* PrkA protein do not seem to play a role in sensing MurA level.

### Effect of mutated MurA variants on ReoM phosphorylation

As our results contradict with the concept of a negative feedback loop and that accumulation of lipid II inhibits PrkA activity, we considered MurA as a direct effector of ReoM phosphorylation. We reasoned that MurA would lose the ability to interfere with ReoM phosphorylation if it is unable to bind ReoM and therefore tested the effect of overproducing the MurA N197D and MurA S262L variants on ReoM phosphorylation, as both substitutions prevent the interaction of MurA with ReoM (35). Remarkably, overproduction of neither of these two MurA variants prevented phosphorylation of ReoM (Fig. 5A), suggesting that a direct interaction between MurA and ReoM is needed.

**Figure 5:**
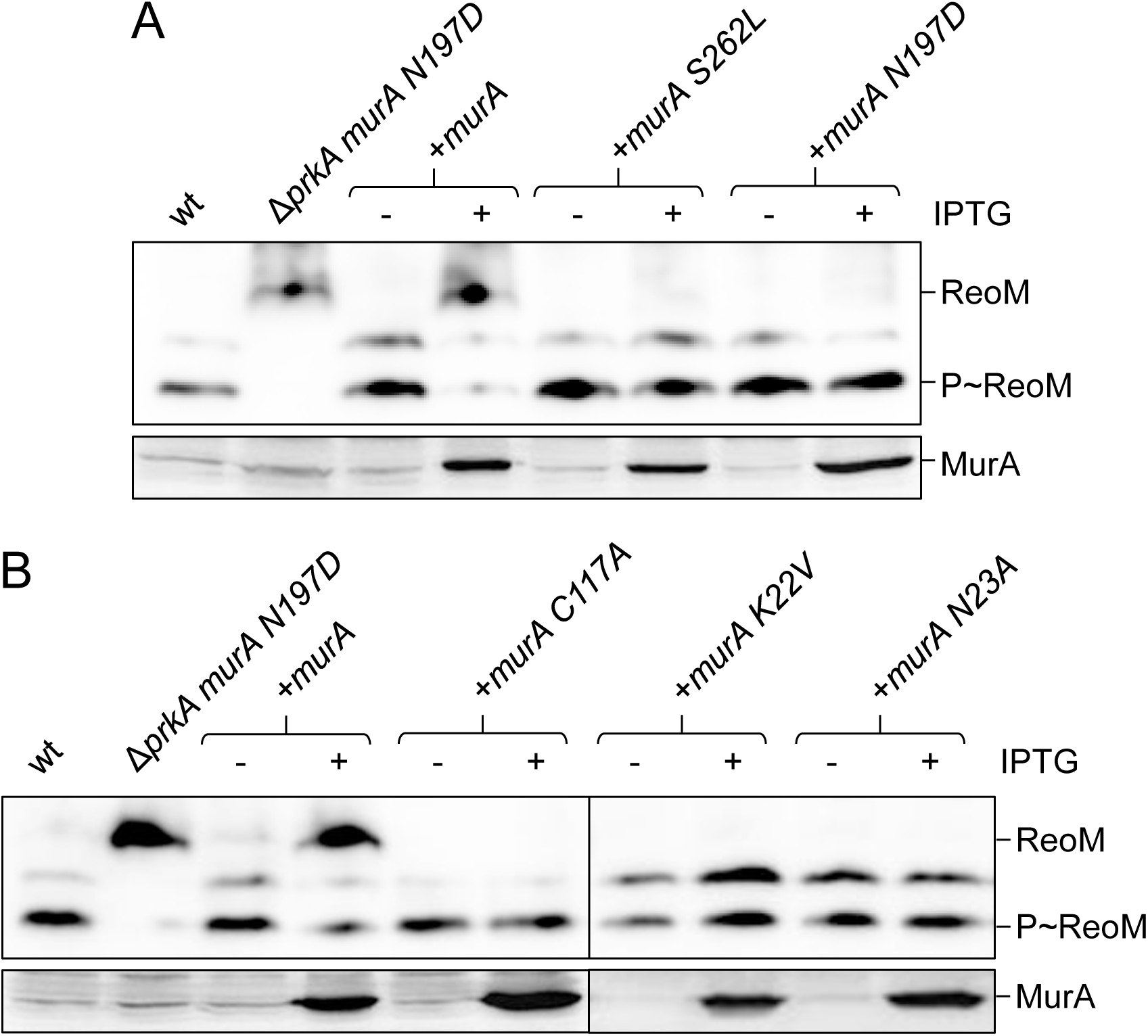
Effect of functional *murA* mutations on ReoM phosphorylation. (A) MurA must interact with ReoM to control its phosphorylation. Western blots showing ReoM phosphorylation (upper blot) and MurA levels (bottom blot) in strains EGD-e (wt), LMS266 (Δ*prkA murA N197D*), LMJR116 (+*murA*), LMSW136 (+*murA S262L*) and LMSW137 (+*murA N197D*) grown in BHI broth ± 1 mM IPTG. (B) The active site cysteine and residues important for substrate binding in MurA are essential for feedback control of ReoM phosphorylation. Western blots showing ReoM phosphorylation (upper blot) and MurA levels (bottom blot) in strains EGD-e (wt), LMS266 (Δ*prkA murA N197D*), LMJR116 (+*murA*), LMPR51 (+*murA C117A*), LMPR57 (+*murA K22V*) and LMPR56 (+*murA N23A*) grown in BHI broth ± 1 mM IPTG.

MurA undergoes substantial conformational changes during substrate binding and catalysis. Unliganded apo-MurA is found in an open conformation, in which the flexible activation loop containing the catalytically important cysteine (C117 in *L. monocytogenes* MurA) is moved away from the enzyme. Upon substrate binding, the activation loop closes the UDP-GlcNAc binding site to bring phosphoenolpyruvate (PEP) into the vicinity of UDP-GlcNAc (*i. e.* closed conformation) (40, 56–58). Remarkably, *E. faecalis* IreB interacts with MurAA only, when MurAA is complexed with UDP-GlcNAc and fosfomycin (20), a condition under which *E. coli* MurA adopts the closed conformation (56).

We tested the effect of inactivating MurA substitutions either preventing UDP-GlcNAc binding (N23A) (59) or progression through the enzymatic cycle (K22V, C117A) (60, 61) on ReoM phosphorylation. This showed that all mutated proteins were unable to suppress phosphorylation of ReoM (Fig. 5B). The N23A protein is supposedly locked in the open conformation, since it cannot bind UDP-GlcNAc (59). The K22V is also catalytically inactive, but not impaired in substrate binding (60). The catalytic cysteine mutant still binds UDP-GlcNAc (62) and then adopts a closed conformation but does not proceed through catalysis (58). This shows that there is no MurA-dependent negative feedback on ReoM phosphorylation when MurA exists in its open unliganded state. Furthermore, MurA must be able to progress through the catalytic cycle to exert this effect.

### The MurA:ReoM interaction is sensitive to ReoM phosphorylation

According to our current model, ReoM phosphorylation regulates the interaction of ReoM with MurA. In this model, only unphosphorylated ReoM can form a complex with MurA and phosphorylation of ReoM prevents this (19, 35). However, this model has mostly been deduced from genetic data and direct evidence for the *L. monocytogenes* proteins is still missing. To test whether the interaction of MurA with ReoM depends on ReoM phosphorylation, we made use of the observation that MurA can be copurified from total cellular extracts with ReoM-His as the bait (35). The essentiality of *prkA* is lost in the absence of *murZ* (35). We therefore generated an isogenic pair of Δ*murZ* strains both expressing *reoM-his* and containing *prkA* (LMPR25) or not (LMPR29). As anticipated, MurA levels in these strains exceeded that of wild type approximately 9-10-fold due to impaired MurA degradation (Fig. 6A, upper and lower panel), regardless whether ReoM was phosphorylated or unphosphorylated (Fig. 6A, middle panel). MurA copurified with ReoM-His from lysates of the *prkA*^+^ strain (Fig. 6B, upper panel) where most of it was phosphorylated (Fig. 6A middle panel). However, the amount of MurA that copurified with ReoM-His from lysates of the Δ*prkA* strain, where ReoM is not phoshproylated, was 3±1-fold higher (n=3, *P*<0.05) albeit less ReoM-His material (0.7±0.1-fold) was pulled down. This is in agreement with the idea that the MurA:ReoM interaction is prevented by phosphorylation of ReoM.

**Figure 6:**
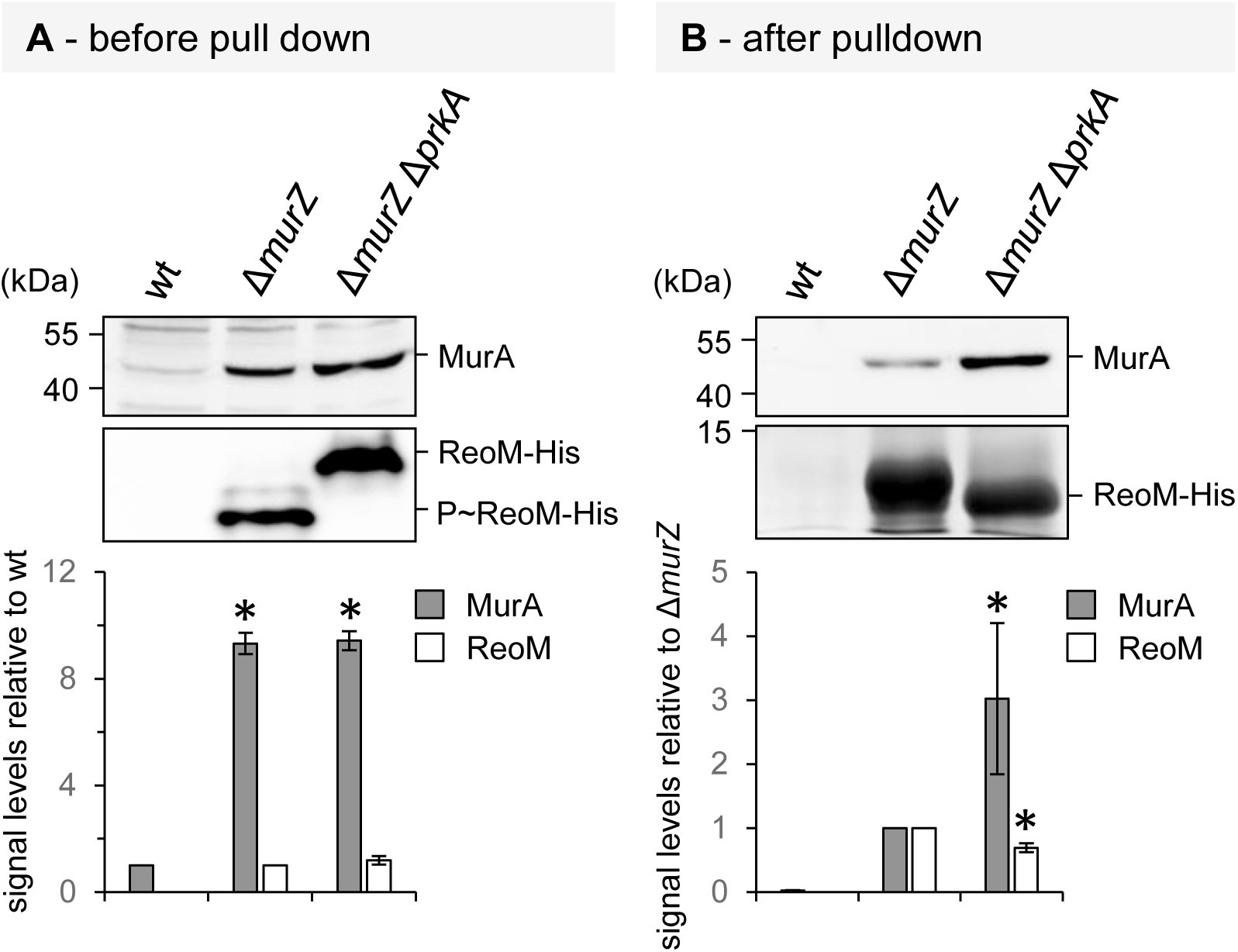
Effect of ReoM phosphorylation on the interaction of ReoM with MurA. Pull down experiment after *in vivo* formaldehyde crosslinking using the *L. monocytogenes* strains EGD-e (wt, control), LMPR25 (Δ*murZ reoM-his*) and LMPR29 (Δ*murZ* Δ*prkA reoM-his*). (A) MurA levels (upper panel) and ReoM phosphorylation (middle panel) prior to crosslinking as determined by Western blotting. For ReoM, samples were separated by native PAGE to allow separation of ReoM species. Signal intensities were quantified by densitometry and expressed as values relative to wild type (lower panel). Average values and standard deviations are shown. Statistically significant differences are labelled by asterisks (*P*<0.01, *t*-test with Bonferroni-Holm correction, n=3). (B) Western blot showing MurA levels (upper panel) and SDS-PAGE showing ReoM levels (middle panel) after pull down. MurA and ReoM amounts were quantified (lower panel) as described above, but expressed relative to the Δ*murZ* sample. Asterisks indicate significance levels (*P*<0.05 for MurA, *P*<0.01 for ReoM, *t*-test with Bonferroni-Holm correction, n=3).

Lastly, we tested whether ReoM would affect MurA activity. This idea was brought forward by Tsui and coworkers, who suggested that unphosphorylated *S. pneumoniae* ReoM could inhibit the activity of the MurA enzymes (31). However, *in vitro* activity of *L. monocytogenes* MurA was not affected by addition of unphosphorylated ReoM (Fig. S6). Thus, the main role of *L. monocytogenes* ReoM is regulation of MurA degradation by ClpCP.

## Discussion

### PASTA kinase activation by GpsB

We demonstrate that *L. monocytogenes* PrkA requires GpsB for full activity. This role of GpsB is specific as none of the other tested cell division proteins contributed to kinase activation and even in non-dividing PBP B1-depleted cells, normal ReoM phosphorylation was observed. Thus, PrkA activity is not dependent on cell division in *L. monocytogenes*, even though septal localization of PASTA kinases in other species suggested such a link (44, 45, 53, 63). Given the phenotypic similarity of *L. monocytogenes gpsB* and *prkA* mutants and suppression of their phenotypes by an almost identical set of mutations (19, 30, 33), the effect of GpsB on PrkA activity was expected. In good agreement, GpsB supported the activity of the *B. subtilis* PrkC kinase *in vitro* and was required for *E. faecalis* IreK activity *in vivo* (42, 43). GpsB belongs to the DivIVA/GpsB family of coiled-coil membrane binding proteins that consist of N-terminal lipid binding domains fused to C-terminal multimerization domains (51). These proteins recruit interacting proteins to curved membrane areas, to which they bind themselves, and partner recruitment can occur either through their N- or C-terminal domain (51, 64). Highly conserved regions in the N-terminal domain of GpsB contribute to membrane and PBP A1 binding and mutations in these regions usually inactivate GpsB (24, 25). We here show that GpsB did not need to bind membranes or to PBP A1 for PrkA activation. However, mutations that abolished the tertiary structure of the hexameric GpsB C-terminus impaired activation of PrkA. This indicates that the C-terminal domain of GpsB plays a prominent role in PrkA activation. Possibly, PrkA interacts with the GpsB C-terminus and as long as the hexameric GpsB structure is maintained, kinase activation can take place. If so, then PrkA would be the first GspB-binding protein described that interacts with the C-terminal domain of GpsB.

### Control of PrkA activity during growth

Control of PASTA kinase activity during cell growth and division is still an unsolved mystery. PASTA kinases are activated by muropeptides and/or lipid II binding to their PASTA domains (44–46). This idea is based on the observation that the PASTA kinases of *S. aureus* and *M. tuberculosis* interact with lipid II *in vitro* and loose this ability when their PASTA domains are deleted or mutated (44, 46). Furthermore, dimerization and septal localization of PASTA kinases is also lost upon deletion of their PASTA domains (44, 53, 63, 65, 66). PASTA domains are not required for kinase activity in general (19, 53, 63, 65), but they are needed for full kinase activity (67). In good agreement, ReoM phosphorylation was also reduced in a *prkA*ΔC mutant (Fig. 2A, Fig. 4D). These findings suggested that lipid II/muropeptides activate PASTA kinases by facilitating their dimerization. However, when lipid II binding residues in the PASTA domains of *M. tuberculosis* PknB were mutated, the protein became hyperactive instead of inactive, rather suggesting inhibition than activation of the kinase activity by lipid II (46). In contrast to this, lipid II activated *S. aureus* PknB *in vitro*, but the degree of activation was only slight (44). Moreover, a structural rather than a regulatory role has recently been proposed for the PASTA domains in *S. pneumoniae* StkP. Through its distal PASTA domain, StkP interacts with the PG hydrolase LytE and positioning of LytE in a defined distance from the membrane, which is determined by the number of PASTA domains, controls cell wall thickness. Obviously, PASTA domains also serve as molecular ruler (63) and PASTA kinase activation models in different bacterial species have not reached convergence.

Our results support the idea that *L. monocytogenes* PrkA is constitutively active. Under most conditions and genetic constellations tested, we observed almost complete ReoM phosphorylation. ReoM is a dimer (19, 68), and therefore existed as ReoM/P∼ReoM hetero- or P∼ReoM/P∼ReoM homodimer, which were the only two ReoM species detected when PrkA was active. Formation of P∼ReoM/P∼ReoM was reduced during stationary phase and ReoM/P∼ReoM as well as unphosphorylated ReoM accumulated instead. Importantly, this growth curve-dependent kinetic required PrpC, which demonstrates that PrpC provides a critical regulatory input and is involved in the shutdown of PG biosynthesis during the transition into stationary phase (Fig. 7).

**Figure 7:**
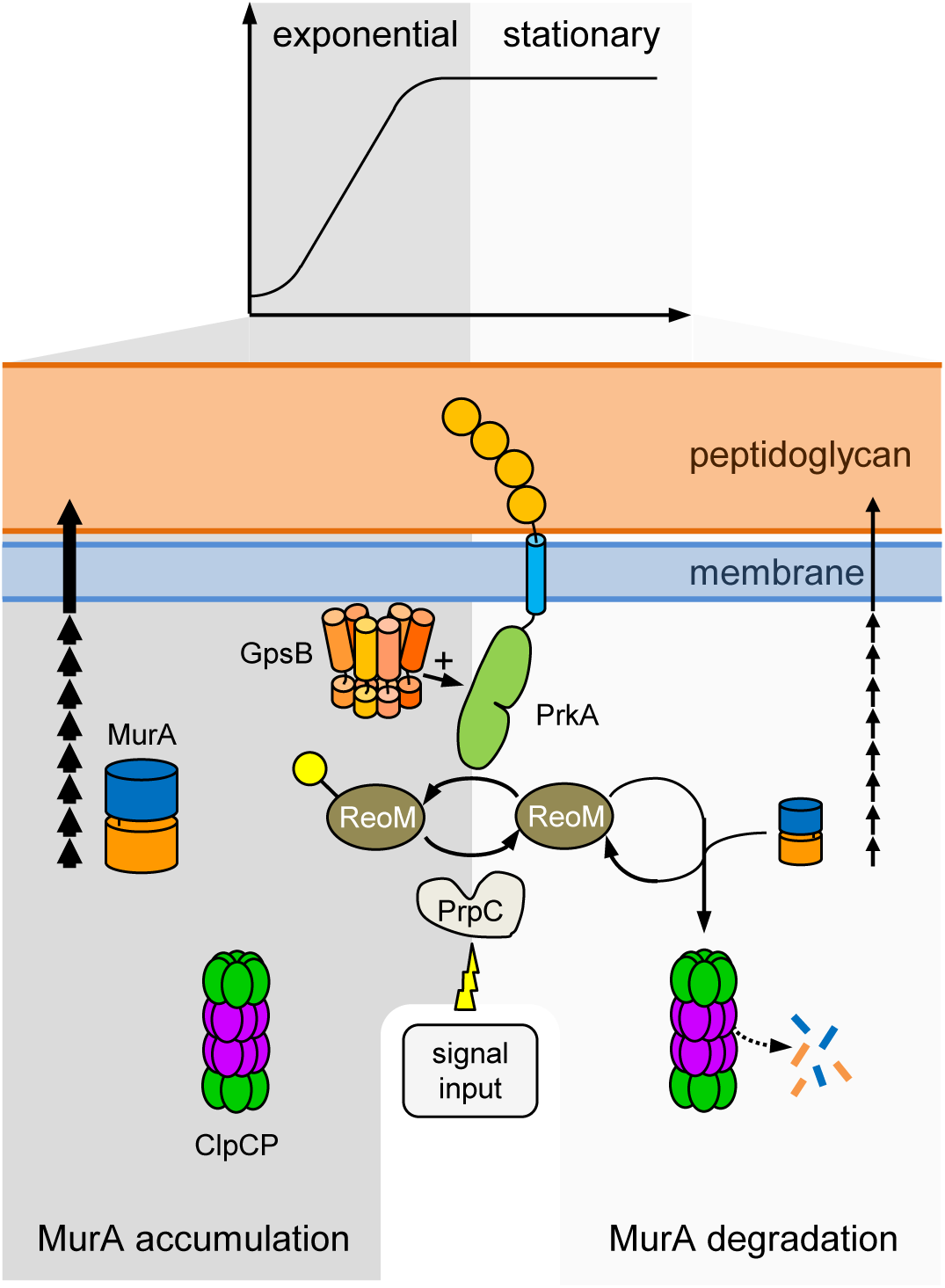
Control of MurA degradation through the phosphorylation status of ReoM. Model describing the known regulators and regulatory influences controlling phosphorylation of ReoM and MurA accumulation in *L. monocytogenes*. During active growth, GpsB-activated PrkA phosphorylates ReoM. Since P∼ReoM cannot interact with MurA, MurA is not degraded and accumulates to ensure PG biosynthesis (left). During stationary phase, P∼ReoM is dephosphorylated, likely due to activation of PrpC. Dephosphorylated ReoM interacts with MurA and transfers MurA to ClpCP for degradation. ReoM itself is not a Clp substrate and is recycled. The transfer of MurA from ReoM to ClpCP also requires ReoY and MurZ, which we have not included in the model for more clarity.

### MurA levels provide a negative feed back on ReoM phosphorylation

A key result of our study was the observation that ReoM phosphorylation is suppressed in mutants that cannot degrade MurA, *i. e.* in Δ*clpC*, Δ*murZ* or Δ*reoY* mutants. Accumulation of MurA in these mutants was the sole reason for this effect, as *murA* overexpression was sufficient to prevent ReoM phosphorylation. Although overexpression of MurA represents an artificial situation, these results imply that there is a negative feedback mechanism linking the level of MurA to ReoM phosphorylation. Inhibition of PrkA by accumulating lipid II would be a fascinating possibility to explain these observations. In such a scenario, spare MurA molecules generating excess lipid II would be proteolytically degraded due to lipid II-inhibited PrkA that leaves ReoM unphosphorylated. However, our results speak against the existence of a such a lipid II-dependent negative feedback loop for three reasons: First, ReoM phosphorylation is still prevented by MurA overexpression in a *prkA*ΔC mutant lacking the PASTA domains. Thus, lipid II binding to PASTA domains cannot be involved. Second, depletion of MurJ activity from Δ*clpC* cells, in which ReoM phosphorylation is prevented due to high MurA levels, does not restore normal ReoM phosphorylation, whereas depletion of MurA from Δ*clpC* cells does. This shows that the signal triggered by excess MurA is generated in the cytoplasmic part of PG biosynthesis somewhere between MurA and MurJ. Third, ReoM phosphorylation depends on the ability of MurA to interact with ReoM. This suggests that MurA itself, its amount or conformation, is possibly sensed by ReoM. MurA also must be catalytically active and able to adopt its closed, substrate-bound conformation to prevent ReoM phosphorylation. This is in good agreement with the results reported by Mascari and coworkers, who showed that substrate binding promotes the interaction of *E. faecalis* MurAA with unphosphorylated IreB *in vitro*, whereas phosphorylated IreB could not bind MurA (20). Our *in vivo* results are consistent with this concept, as copurification of MurA with ReoM was sensitive to ReoM phosphorylation. Most of our findings would be compatible with a ReoM partner switching model (Fig. 7). Considering the observations that unphosphorylated ReoM forms complexes with PrkA (19) and MurA (20, 35), it seems conceivable that PrkA and MurA compete for ReoM as their shared interaction partner. If the amount of MurA is low, ReoM preferably interacts with PrkA leading to ReoM phosphorylation. When the MurA level rises, complex formation of ReoM with MurA becomes more likely and this would initiate MurA degradation. In this sense, would function as a security valve that ensures adaptation of the MurA level to its own amount. In addition, the ReoM phosphorylation equilibrium responds to alterations in PrpC activity and this signal perception route ensures that PG biosynthesis can be shut down during stationary phase (Fig. 7). Furthermore, saturation of MurA with its substrates is integrated into the control network (Fig.5B) (20), but the significance of this is not yet clear.

The partner switching model predicts that ReoM phosphorylation is enhanced in a *murA N197D* mutant because MurA N197D has lower ReoM binding affinity (35). However, we did not observe this (Fig. 2A), suggesting that the N197D mutation has a more complex effect on the ReoM phosphorylation equilibrium than previously thought or that the model needs to be extended. The *in vitro* reconstitution of this system combined with mathematical modelling of the kinetic parameters could provide an even better understanding of the control of ReoM phosphorylation in the future.

## Materials and Methods

### Bacterial strains and growth conditions

Tab. 1 lists all bacterial strains and plasmids used in this study. *L. monocytogenes* strains were generally grown in BHI broth or on BHI agar plates at 37°C. Erythromycin (5 µg ml^-1^), kanamycin (50 µg ml^-1^), X-Gal (100 µg ml^-1^) or IPTG (1 mM) were added as indicated. *Escherichia coli* TOP10 was used as the standard cloning host (69).

**Table 1:**
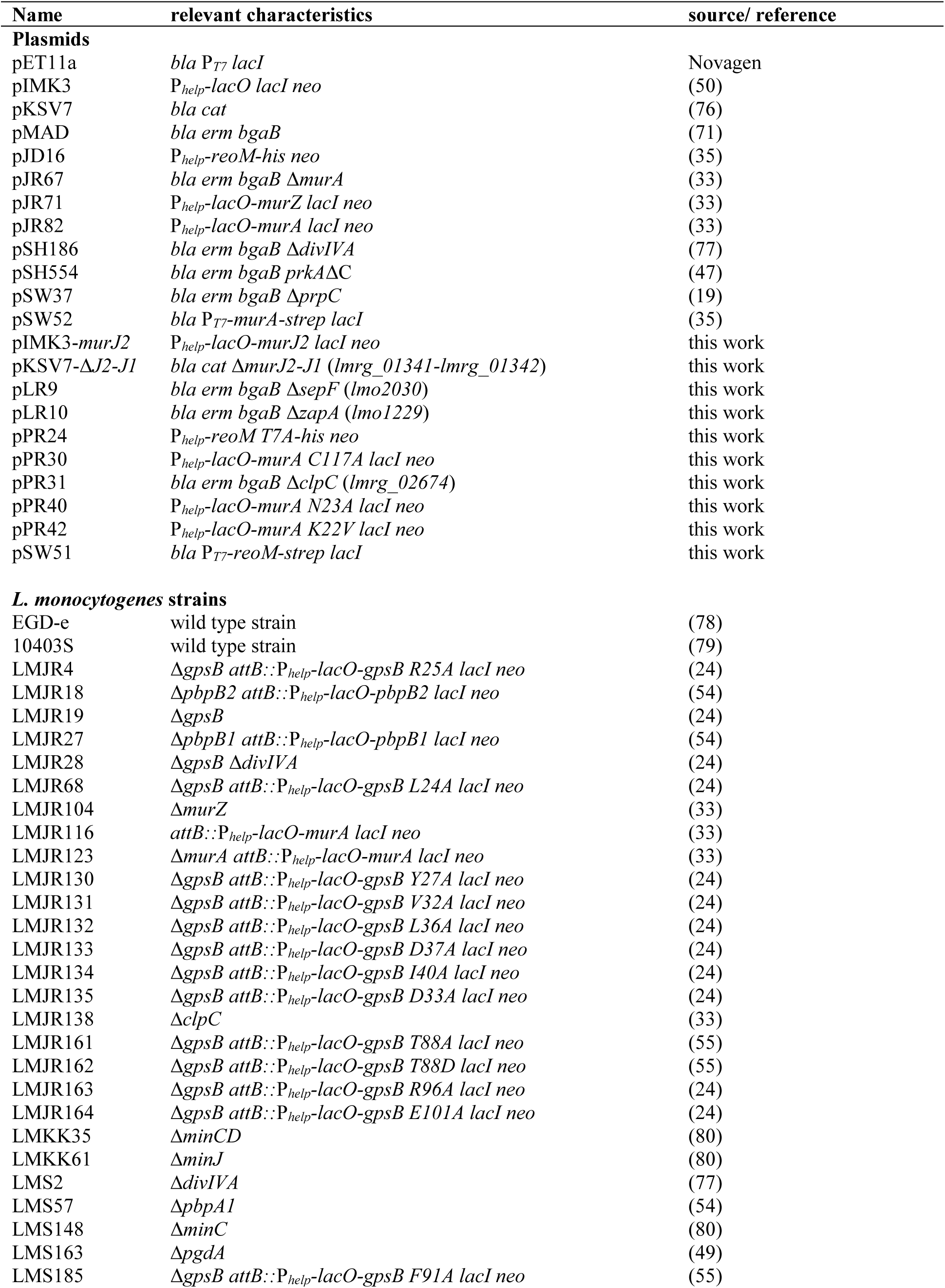
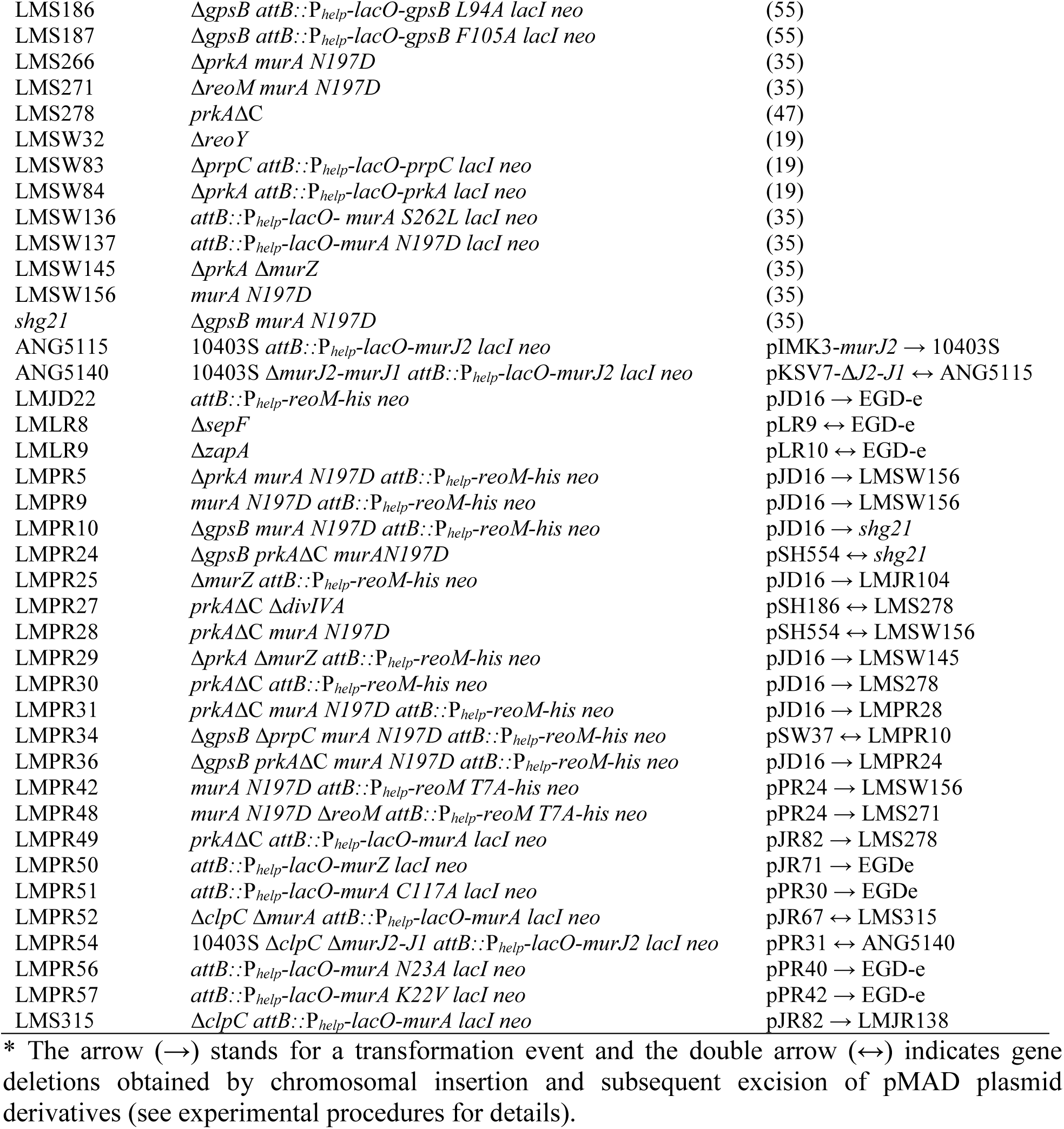
Plasmids and strains used in this study.

### General methods, manipulation of DNA and oligonucleotide primers

Standard methods were used for transformation of *E. coli* and for isolation of plasmid DNA (69). Transformation of *L. monocytogenes* was performed using electroporation as described by others (50). PCR, restriction and ligation of DNA was performed following the manufactureŕs instructions. All primer sequences are listed in Tab. 2. Ceftriaxone minimal inhibitory concentrations were determined as described previously using E-test strips with a concentration range of 0.016–256 μg/ml (Bestbiondx, Germany) (54). Resistance against lysozyme was determined in an autolysis assay as described previously (49). Minimal inhibitory concentrations of lysozyme were determined as described (70).

**Table 2:**
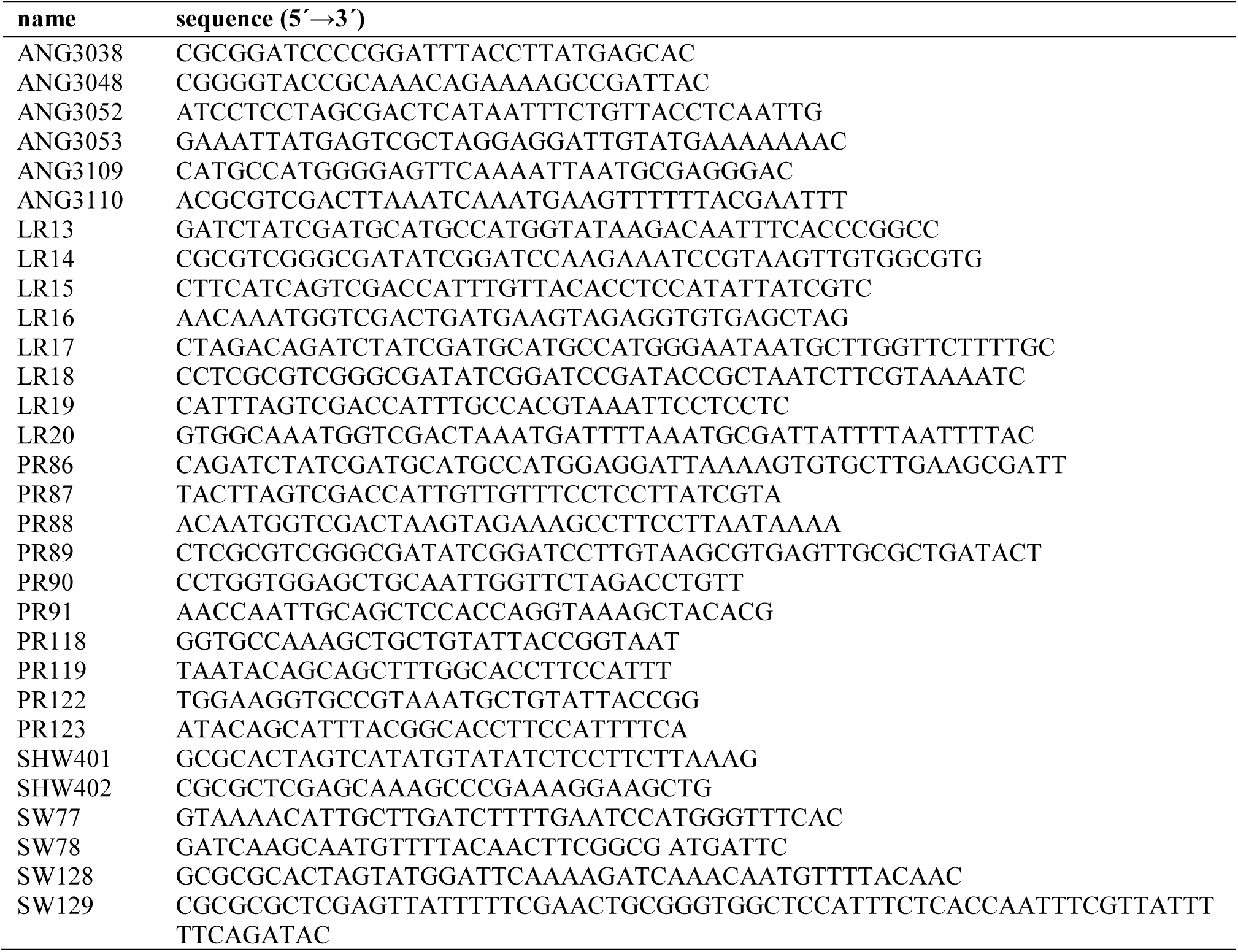
Primers used in this study.

### Microscopy

Cell membranes were stained by addition of 1 μl nile red solution (100 μg/ml in DMSO) to 100 μl of exponentially growing bacteria. Images were taken with a Nikon Eclipse Ti microscope coupled to a Nikon DS-MBWc CCD camera and processed using the NIS elements AR software package (Nikon). Cell lengths were measured using the distance measuring tools of NIS elements AR.

### Construction of plasmids and strains

Plasmid pSW51 was constructed for overexpression of ReoM-Strep in *E. coli*. To this end, *reoM* was amplified from chromosomal DNA using SW128/SW129 as the primers. The resulting DNA fragment was cut using SpeI/XhoI and then ligated with the backbone of plasmid pET11a, which had been linearized in a PCR using the oligonucleotides SHW401/SHW402 (introducing the recognition sites for SpeI/XhoI) and cut with the same enzymes.

Plasmid pJR24 was obtained by introducing the T7A mutation into the *reoM* gene of plasmid pJD16 by quikchange mutagenesis and oligonucleotides SW77/SW78 as the primers.

For deletion of *sepF*, plasmid pLR9 was constructed. Up- and downstream fragments of *sepF* were amplified in a PCR using LR13/LR15 and LR16/LR14, respectively, as the primers. Both fragments were fused together by SOE-PCR and the resulting fragment was inserted into pMAD by restriction free cloning.

Plasmid pLR10, constructed for deletion of *zapA*, was obtained in a similar way. Here, up- and downstream fragments of *zapA* were amplified in a PCR using the oligonucleotides LR17/LR19 and LR18/LR20, respectively. The two fragments were fused together by SOE-PCR and then inserted into pMAD by restriction-free cloning.

Plasmid pPR30 was constructed for expression of an enzymatically inactive *murA* variant. For this purpose, the C117A mutation was introduced into the *murA* gene present on plasmid pJR82 by quikchange mutagenesis with PR90 and PR91 as the mutagenic primers. Likewise, the K22V and N23A mutations were introduced into *murA* of pJR82 using primers PR122/PR123 (yielding pPR42) and PR118/PR119 (for pPR40), respectively.

Plasmid pPR31 was constructed for deletion of *clpC* from the 10403S chromosome. To this end, regions up- and downstream of *clpC* (*lmrg_02674*) were amplified from 10403S chromosomal DNA using the primer pairs PR86/PR87 and PR88/89 and spliced together in a SOE-PCR. The resulting fragment was then cloned into pMAD using BamHI/NcoI.

For IPTG-dependent expression of *murJ2*, the *murJ2* open reading frame was amplified from 10403S chromosomal DNA using the primer pair ANG3109/ANG3110, the resulting PCR fragment was cut with NcoI and SalI and ligated with pIMK3 that had been cut with the same enzymes.

For the markerless in-frame deletion of *murJ1* and *murJ2*, 1-kb DNA-fragments upstream of *murJ2* and downstream of *murJ1* were amplified by PCR using primer pairs ANG3038/ANG3052 and ANG3053/ANG3048, respectively. The resulting PCR products were fused in a second PCR using primers ANG3038/ANG3048, cut with BamHI and KpnI and ligated with plasmid pKSV7 that had been cut with the same enzymes.

pIMK3 derivatives were brought into *L. monocytogenes* strains by electroporation and transformants were selected on BHI agar plates containing kanamycin at 37°C. Plasmid integration at the tRNA^Arg^ *attB* site was confirmed by PCR. Likewise, pMAD derivatives were introduced into *L. monocytogenes*, but selection was carried out on BHI agar plates containing X-Gal and erythromycin at 30°C. The plasmid integration-excision protocol described by Arnaud *et al.* (71) was used for gene deletions in the EGD-e background. Plasmid pKSV7-Δ*murJ2-J1* was brought into *L. monocytogenes* 10403S strains by electroporation. The protocol of Camilli *et al.* (72) was used for the construction of the *murJ2-J1* deletion strain by allelic exchange. All gene deletions were confirmed by PCR.

### Protein purification and antibody generation

ReoM-Strep was overexpressed in *E. coli* BL21, which was cultivated in LB broth containing ampicillin (100 µg/ml) at 37°C. Expression of ReoM-Strep was induced at an optical density of OD_600_=0.5 by addition of 1 mM IPTG (final concentration). The culture was grown over night at 18°C before cells were harvested by centrifugation. The cell pellet was washed once with ZAP buffer (10 mM Tris/HCl pH 7.5, 200 mM NaCl), and disrupted in ZAP buffer containing 1 mM PMSF using an EmulsiFlex homogeniser (Avestin, Germany). Cell debris was removed by centrifugation (6,000 *g*, 5 min, 4°C) and the resulting supernatant was ultracentrifuged (60,000 *g*, 30 min 4°C) to remove any remaining particles. ReoM-Strep was purified using affinity chromatography and Strep-Tactin® Sepharose (IBA Lifesciences, Germany) according to the manufacturer’s instructions. Eluted fractions containing ReoM-Strep were pooled and the buffer was exchanged against PBS using PD10 desalting columns. Samples were aliquoted and stored at −20°C. MurA-Strep was purified from *E. coli* BL21 carrying plasmid pSW52 as described previously (35). ReoM-Strep was used to immunize a rabbit for the generation of a polyclonal antiserum and the IgG fraction was purified from the serum (Biogenes, Germany).

### MurA activity assay

MurA activity was determined by measuring the amount of phosphate released from PEP. For this purpose, 2.5 μg of MurA were mixed with 10 mM uridine 5′-diphospho-*N*-acetylglucosamine (UDP-GlcNAc, Sigma-Aldrich) in a reaction volume of 50 µl containing 100 mM Tris/HCl pH8.0 and 150 mM NaCl as the buffer and preincubated at 37°C for 15 min. 5 μl 10 mM phosphoenolpyruvate (Sigma-Aldrich) was added to start the reaction. After 30 min of incubation at room temperature, 800 μl staining solution were added to the reaction. The staining solution was freshly prepared from 10 ml ammonium molybdate solution (4.2 g in 100 ml 4 M HCl), 30 ml malachite green solution (225 mg malachite green in 500 ml H_2_O) and 40 μl Triton X-100. Absorption was measured at λ = 660 nm, corrected for background in the absence of UDP-GlcNAc and used to calculate the amount of phosphate released using a standard curve that was generated using solutions with different phosphate concentrations (0 mM – 0.5 mM Na_2_HPO_4_ in 100 mM Tris/HCl pH8.0, 150 mM NaCl).

### Isolation of cellular proteins

30 ml of BHI broth were inoculated with an overnight culture to an OD_600_ = 0.05 and grown to an OD_600_ of ∼1.0 unless otherwise stated. Cell cultures were harvested by centrifugation, washed with ZAP buffer (50 mM Tris/HCl pH 7.5, 200 mM NaCl), resuspended in 0.5 ml ZAP buffer also containing 1 mM PMSF and disrupted by sonication. Cellular debris were removed by centrifugation and the supernatant was used as total cellular protein extract.

### Native page, SDS-PAGE and Western blotting

To separate proteins on a native gel, 50 µg protein extract were mixed with 6x native loading dye (50 mM Tris/HCl pH 8.5, 0.1 % (w/v) bromophenol blue, 10 % (w/v) glycerol) and separated on a non-denaturing 15% polyacrylamide (PA) gel (in 375 mM Tris/HCl pH8.8 buffer) that was overlaid with a 5% PA collecting gel in 125 mM Tris/HCl pH6.8 as the buffer. Both gels were prepared using a 37.5:1 PA solution. Electrophoresis was performed with running buffer (25 mM Tris/HCl pH 8.8, 129 M glycine) at 100 V for 2.5 h at room temperature. For separation under denaturing conditions, protein extracts were separated using standard SDS-PAGE. Transfer of proteins from both types of gels onto positively charged polyvinylidene fluoride membranes was performed using a semi-dry transfer unit. ReoM and MurA were detected using polyclonal rabbit antisera recognizing *L. monocytogenes* ReoM (this work) and *B. subtilis* MurAA (73), respectively, as the primary antibodies. An anti-rabbit immunoglobulin G conjugated to horseradish peroxidase was used as the secondary antibody. Detection of antibody antigen complexes was performed using the ECL chemiluminescence detection system (Thermo Scientific) in a chemiluminescence imager (ChemiDoc MP Imaging System, BioRad).

### *In vivo* formaldehyde cross-linking and pull down

*L. monocytogenes* strains expressing His-tagged bait proteins were cultivated in 500 ml BHI broth containing 1 mM IPTG at 37 °C until an OD_600_ of 1.0 was reached. Cultures were treated with formaldehyde at a final concentration of 1% for 30 min. Cross-linking was quenched by glycine addition (50 mM final concentration) for 5 min. Cells were harvested by centrifugation and resuspended in 2 ml UT buffer (0.1 M HEPES, 0.5 M NaCl, 50 mM imidazole, 8 M urea, 1% Triton X-100, 1 mM PMSF, 1 mM dithiothreitol [DTT]). Cell disruption by sonication was performed for 60 min on ice and cell debris was removed by centrifugation. Cleared protein extracts were incubated with 200 μl of MagneHis solution (Promega, USA) overnight, rotating at room temperature. The MagneHis particles were then washed five times with 2 ml of UT buffer. Protein complexes were eluted by incubation in 500 µl elution buffer (0.1 M Tris-HCl pH 7.5, 0.5 M imidazole, 1% SDS, 10 mM DTT) at room temperature for 30 min. Eluates were concentrated five-fold using centrifugal micro-concentrators. Samples were mixed with SDS-PAGE loading dye, and cross-linking was reversed by heating at 95°C for 1 h. Decrosslinked samples were separated by SDS-PAGE and analysed by Western blotting.

### Purification of His-tagged ReoM from *L. monocytogenes* under native conditions

For purification of His-tagged ReoM from *L. monocytogenes* cells under native conditions, we followed the same protocol as outlined above for the pull down experiments except that the cross-linking and quenching reactions were skipped.

### In gel digestion for mass spectrometry

Protein bands were prepared for mass spectrometry using the protocol of Shevchenko *et al.* (74). Resulting peptides were desalted using 200 µL StageTips packed with two Empore™ SPE Disks C18 (3M Purification, Inc., Lexington, USA) according to Rappsilber *et al.* (75) and concentrated using a vacuum concentrator. Samples were resuspended in 12 µL 0.1 % formic acid.

### nLC-MS/MS

Peptides were analysed on an EASY-nanoLC 1200 (Thermo Fisher Scientific, Bremen, Germany) coupled online to a Q Exactive™ HF mass spectrometer (Thermo Fisher Scientific, Bremen, Germany). 5 µL sample was injected onto a PepSep column (15 cm length, 75 µm i.d., 1.5 µm C18 beads, PepSep, Marslev, Denmark) using a stepped 60 min gradient of 80% acetonitrile (solvent B) in 0.1% formic acid (solvent A) at 300 nL/min flow rate: 4–8% B in 5:06 min, 8-26% B in 41:12 min, 26–31% B in 6:00 min, 31–39% B in 4:12 min, 39–95% B in 0:10 min, 95% B for 2:20 min, 95–0% B in 0:10 min and 0% B for 0:50 min. Column temperature was kept at 50°C using a butterfly heater (Phoenix S&T, Chester, PA, USA). The Q Exactive™ HF was operated in a data-dependent manner in the m/z range of 300 – 1,650. Full scan spectra were recorded with a resolution of 60,000 using an automatic gain control (AGC) target value of 3 × 10^6^ with a maximum injection time of 100 ms. Up to the 10 most intense 2+-4+ charged ions were selected for higher-energy c-trap dissociation (HCD) with a normalized collision energy (NCE) of 27%. Fragment spectra were recorded at an isolation width of 2 Th and a resolution of 30,000@200m/z using an AGC target value of 1 × 10^5^ with a maximum injection time of 50 ms. The minimum MS² target value was set to 1 × 10^4^. Once fragmented, peaks were dynamically excluded from precursor selection for 30 s within a 10 ppm window. Peptides were ionized using electrospray with a stainless-steel emitter, I.D. 30 µm, (Proxeon, Odense, Denmark) at a spray voltage of 2.1 kV and a heated capillary temperature of 275°C.

### Analysis of mass spectrometric data

The mass spectra were analysed using Proteome Discoverer 2.5 (Thermo Fisher Scientific, Bremen, Germany). Spectra were analysed using SequestHT with a tolerance of 10 ppm in MS^1^ and 0.02 Da in HCD MS² mode, strict trypsin specificity and allowing up to two missed cleavage sites. Cysteine carbamidomethylation was set as a fixed modification and oxidation (M), phosphorylation (S,T,Y) as well as N-terminal acetylation and loss of initial methionine as variable modifications. Peptides were identified at 1 % false discovery rate using Percolator and quantified using Minora feature detector with default settings. The localization of phosphorylation sites was scored using ptmRS. Afterwards phosphorylated peptides were further filtered using the cross-correlation score (Xcorr) (z=2 > 2, z=3 > 2.3, z=4 > 2.6), best site probability (> 0.8) and MS^1^ mass accuracy (< 5 ppm) to retrieve a list of high confident phosphopeptides.

## Acknowledgments

This work was funded by DFG grants HA 6830/1-2 and HA6830/4-1 (to S. H.). The authors would like to thank Janina Döhling for her help with some experiments.

