## Supplementary Figures S1-S6 for "Cytosolic factors controlling PASTA kinase-dependent ReoM phosphorylation"

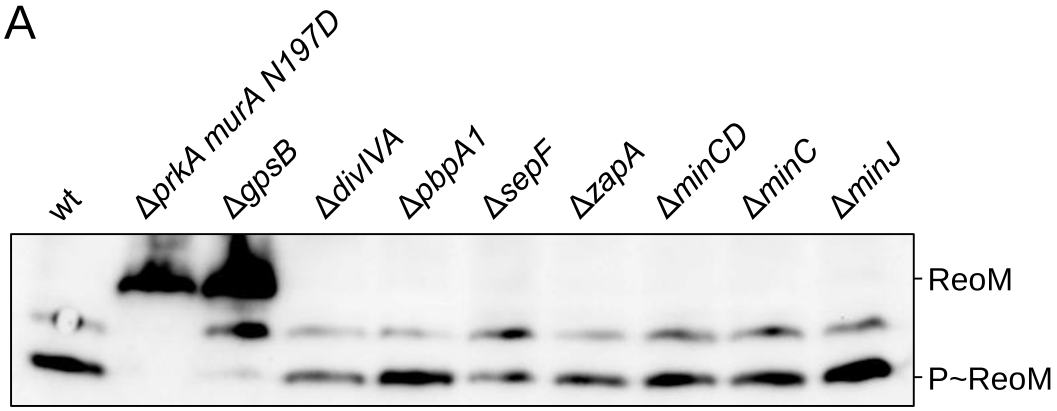


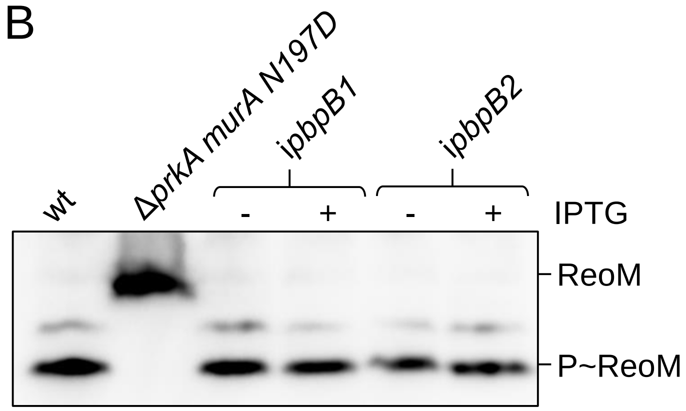


**Figure S1: Influence of cell division genes on ReoM phosphorylation.**

(A) Western blot after native PAGE for analysis of ReoM phosphorylation in *L. monocytogenes* strains lacking non-essential cell division genes. Samples were obtained from *L. monocytogenes* strains EGD-e (wt), LMS266 (Δ*prkA murA N197D*), LMJR19 (Δ*gpsB*), LMS2 (Δ*divIVA*), LMS57 (Δ*pbpA1*), LMLR8 (Δ*sepF*), LMLR9 (Δ*zapA*), LMKK35 (Δ*minCD*), LMS148 (Δ*minC*) and LMKK61 (Δ*minJ*) that were grown in BHI broth at 37°C to an optical density of 1.0.

(B) Effect of PBP B1 and PBP B2 depletion on phosphorylation of endogenous ReoM. Strains EGD-e (wt), LMS266 (Δ*prkA murA N197D*), LMJR27 (i*pbpB1*) and LMJR18 (i*pbpB2*) were grown under the same conditions as above. IPTG was added as indicated.

Rothe *et al.* Fig. S1

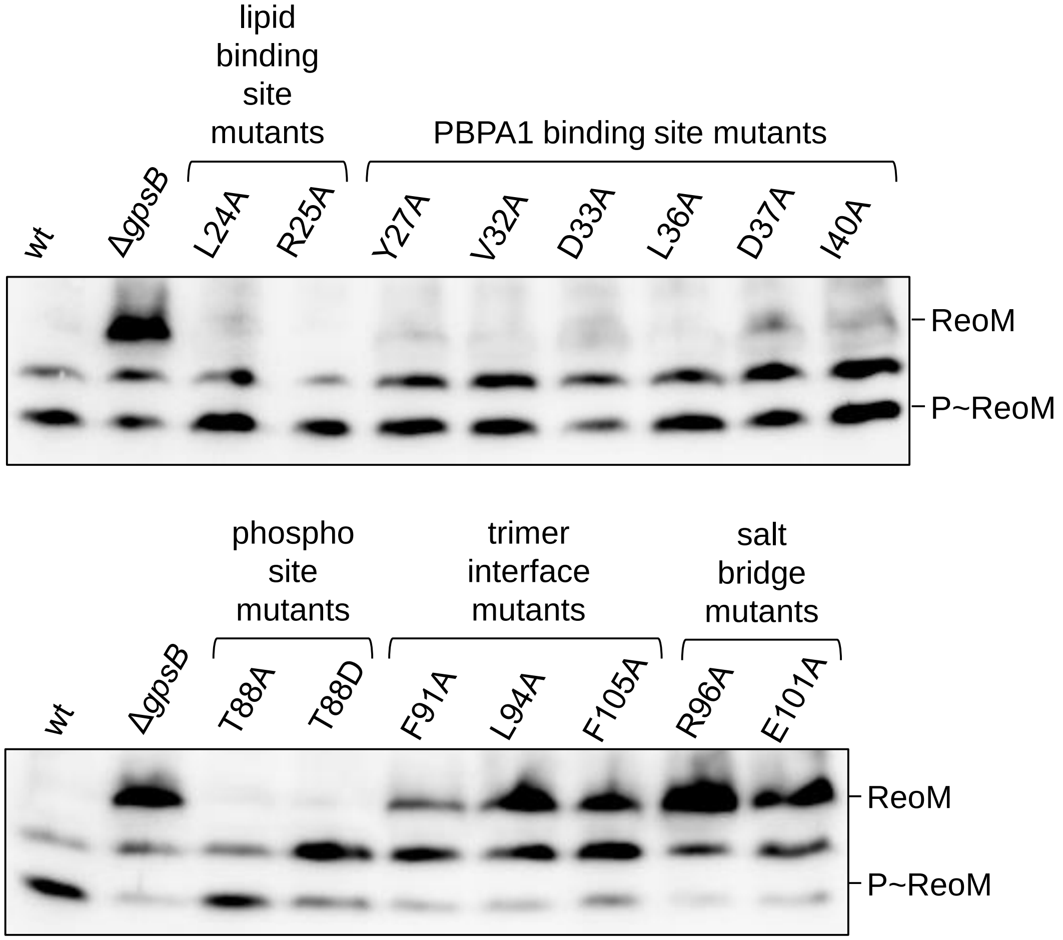


**Figure S2: Effect of functional *gpsB* mutations on PrkA activity.**
Western blot after native PAGE for analysis of *in vivo* ReoM phosphorylation in *L. monocytogenes* strains expressing selected *gpsB* mutant alleles. *L. monocytogenes* strains were EGD-e (wt), LMJR19 (Δ*gpsB*), LMJR68 (L24A), LMJR4 (R25A), LMJR130 (Y27A), LMJR131 (V32A), LMJR135 (D33A), LMJR132 (L36A), LMJR133 (D37A), LMJR134 (I40A), LMJR161 (T88A), LMJR162 (T88D), LMS185 (F91A), LMS186 (L94A), LMS187 (F105A), LMJR163 (R96A) and LMJR164 (E101A)

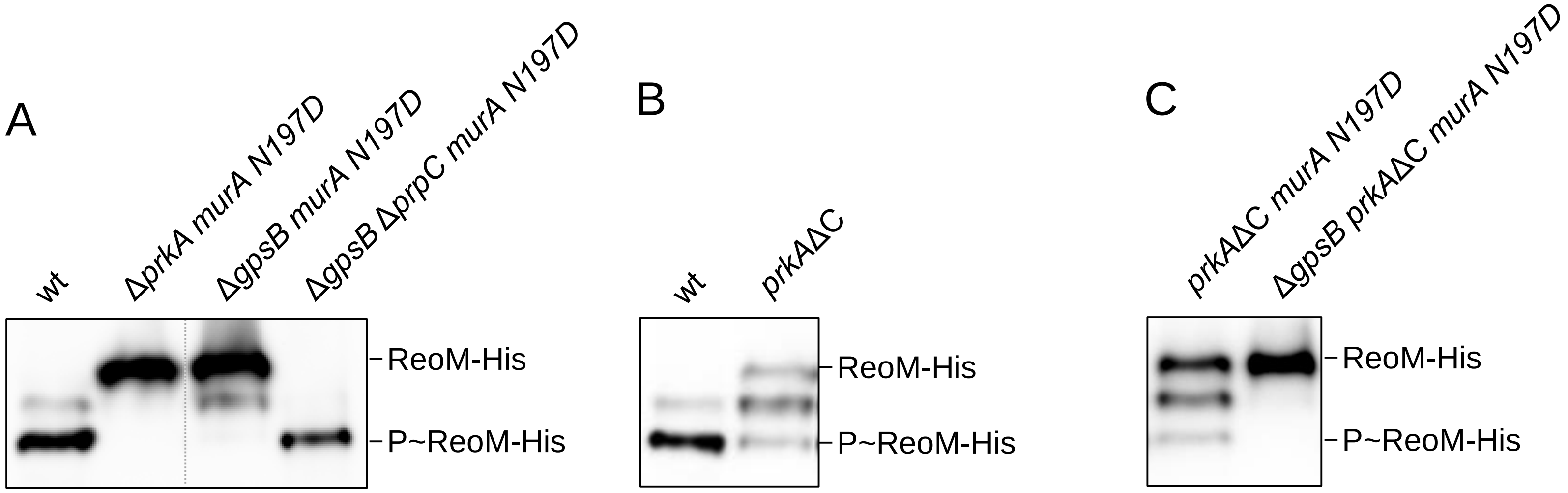


**Figure S3: Role of PrpC and the PrkA PASTA domains in ReoM phosphorylation**

Western blots after native PAGE to separate the differently phosphorylated forms of ReoM in total cellular extracts. Strains were cultivated in BHI broth at 37°C up to an optical density of 1.0.

(A) Effect of PrpC on phosphorylation of ReoM. Samples were from ReoM-His producing *L. monocytogenes* strains LMJD22 (wt), LMPR5 (Δ*prkA murA N197D*), LMPR10 (Δ*gpsB murA N197D*) and LMPR34 (Δ*gpsB* Δ*prpC murA N197D*).

(B) Contribution of the PrkA PASTA domains to ReoM phosphorylation. Strains used were LMJD22 (wt), and LMPR30 (*prkA*ΔC), which all expressed *reoM-his*.

(C) P~ReoM formation in the *prkA*ΔC mutant is still GpsB-dependent. Strains compared with each other here were ReoM-His producing LMPR31 (*prkA*ΔC *murAN197D*) and LMPR36 (Δ*gpsB prkA*ΔC *murAN197D*).

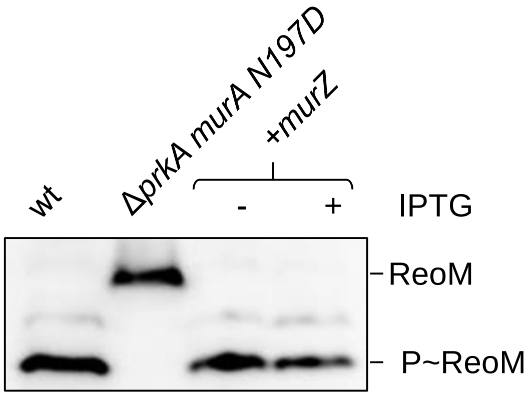


**Figure S4:Effect of MurZ overexpression on ReoM phosphorylation.**
Western blot showing ReoM phosphorylation in strains EGD e (wt), LMS266 (Δ*prkA murA N197D*) and LMPR50 (+*murZ*) grown in BHI broth ± 1 mM IPTG.

Rothe *et al.* Fig. S4

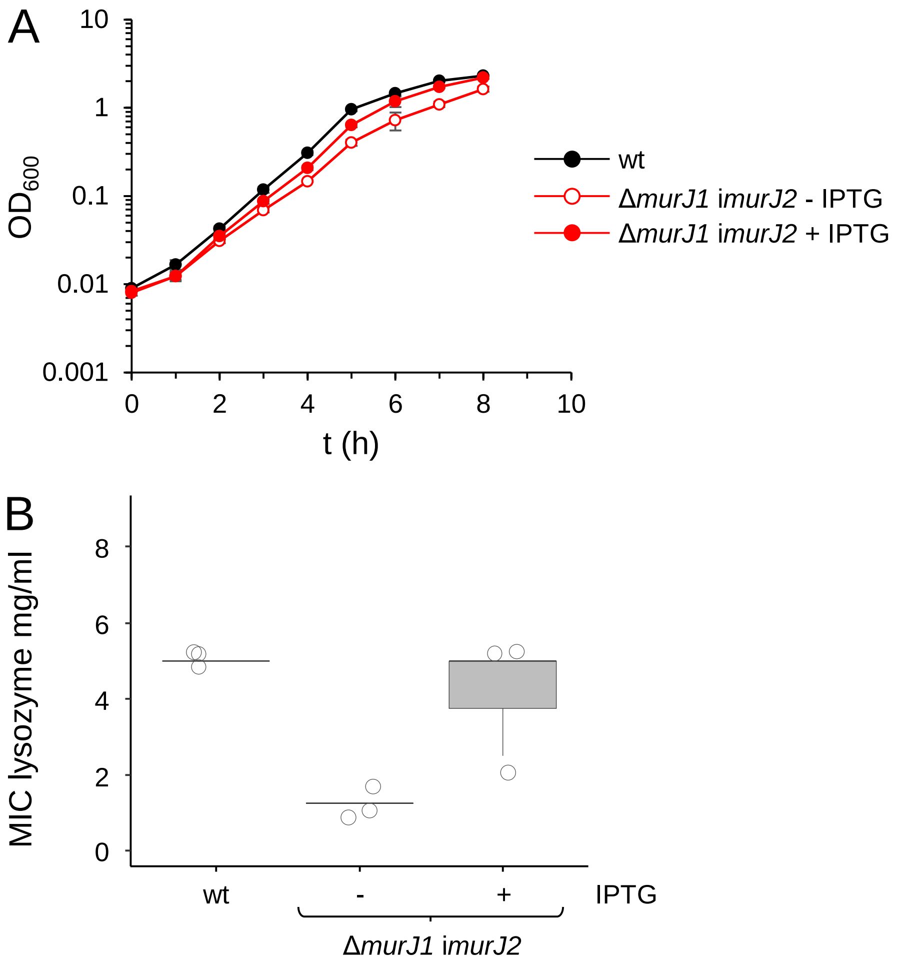


**Figure S5: Depletion of MurJ activity**
(A) Effect of MurJ depletion on growth of *L. monocytogenes*. Growth of *L. monocytogenes* strains 10403S (wt) and ANG5140 (Δ*murJ1 imurJ2*) in BHI broth ± 1 mM IPTG at 37°C. Pre-depleted precultures were used as inoculum. The experiment was repeated three times and average values and standard deviations are shown.
(B) Effect of MurJ depletion on lysozyme resistance. Minimal inhibitory concentrations of lysozyme were determined in a growth experiment, in which the same strains as in panel A were grown in BHI broth ± 1 mM IPTG at 37°C in the presence of increasing lysozyme concentrations. Data collected from three independent experiments are shown.

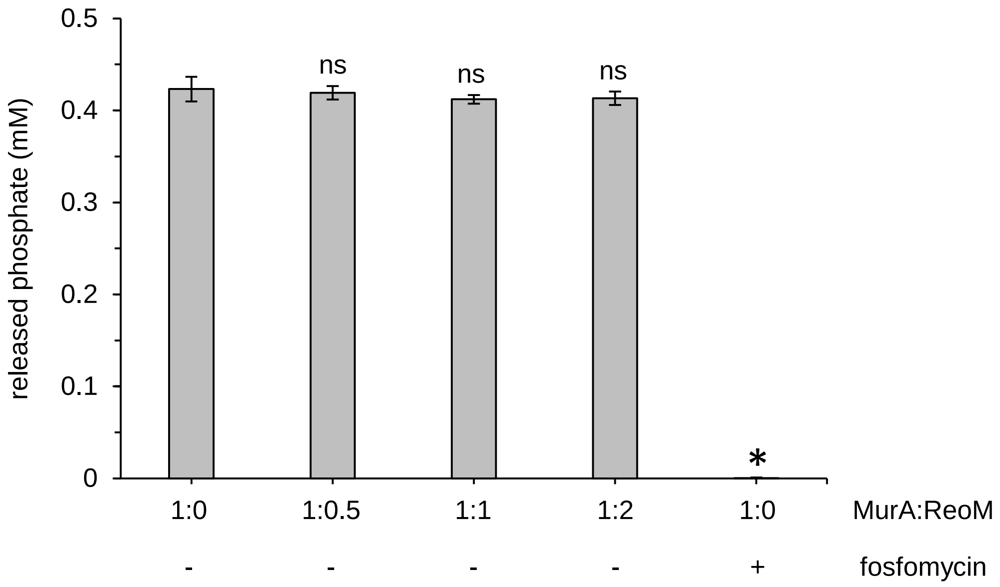


**Figure S6: Effect of unphosphorylated ReoM on MurA activity.**
*In vitro* activity of *L. monocytogenes* MurA in the presence of different ReoM concentrations. MurA activity was measured through the determination of the amount of released phosphate after 30 min of incubation in the presence of UDP-GlcNAc and PEP. Average values and standard deviations were calculated from technical replicates (n=3). Levels of statistical significance are shown (* $P$<0.01, ns – not significant, *t*-test with Bonferroni-Holm correction).

Rothe *et al.* Fig. S6
